# Time-resolved systems analysis reveals a critical role of XCR1+ dendritic cells in the maintenance of effector T cells during chronic viral infection

**DOI:** 10.1101/476077

**Authors:** Jordi Argilaguet, Mireia Pedragosa, Anna Esteve-Codina, Graciela Riera, Enric Vidal, Cristina Peligero-Cruz, David Andreu, Tsuneyasu Kaisho, Gennady Bocharov, Burkhard Ludewig, Simon Heath, Andreas Meyerhans

**Affiliations:** Infection Biology Laboratory, Department of Experimental and Health Sciences (DCEXS), Universitat Pompeu Fabra, Barcelona, Catalonia 08003, Spain.; CNAG-CRG, Center for Genomic Regulation (CRG), Barcelona Institute of Science and Technology, 08028 Barcelona, Spain.; Universitat Pompeu Fabra (UPF), Barcelona, Catalonia 08003, Spain; IRTA, Centre de Recerca en Sanitat Animal (CReSA-IRTA-UAB), Campus de la Universitat Autònoma de Barcelona, 08193 Bellaterra, Barcelona, Catalonia, Spain.; Laboratory of Proteomics and Protein Chemistry, DCEXS, Universitat Pompeu Fabra, 08003 Barcelona, Spain.; Department of Immunology, Institute of Advanced Medicine, Wakayama Medical University, Wakayama 641-8509, Japan.; Laboratory for Immune Regulation, World Premier International Research Center Initiative, Immunology Frontier Research Center, Osaka University, Osaka 565-0871, Japan.; Institute of Numerical Mathematics, Russian Academy of Sciences, Moscow 119333, Russia.; Institute for Immunobiology, Kantonsspital St. Gallen, 9007 St. Gallen, Switzerland.; Institució Catalana de Recerca i Estudis Avançats (ICREA), Barcelona, 08003, Spain.

## Abstract

Upon a viral infection, the host immune system attempts to eradicate the virus. However, once the infection threat seems overwhelming, the infected host actively shuts down effector responses to reduce immunopathology. The price to pay for this is the establishment of a chronic infection that is only partially controlled by a lower level immune response. The genetic networks underlying this infection fate decision and the immune adaptation to the lower level response are not well understood. Here we used an integrated approach of gene coexpression network analysis of time-resolved splenic transcriptomes and immunological analysis to characterize the host response to acute and chronic lymphocytic choriomeningitis virus (LCMV) infections. We found first, an early attenuation of inflammatory monocyte/macrophage prior to the onset of T cell exhaustion and second, a critical role of the XCL1-XCR1 communication axis during the functional adaptation of the T cell response to the chronic infection state. These findings not only reveal an important feedback mechanism that couples T cell exhaustion with the maintenance of a lower level of effector T cell response but also suggest therapy options to better control virus levels during the chronic infection phase.

**Author Summary:** The outcomes of viral infections are the result of dynamic interplays between infecting viruses and induced host responses. They can be categorized as either acute or chronic depending on temporal virus-host relationships. Chronic infections are associated with immune exhaustion, a partial shut-down of effector responses. The processes underlying infection fate decisions are incompletely understood. Here we analyzed, on a systems level, infection-fate–specific gene signatures and the resulting adaptive processes of the host. We used the well-established lymphocytic choriomeningitis virus infection mouse model which has been instrumental to detect many fundamental processes in the virus-immune system crosstalk that are also relevant in human infections. We show an early attenuation of macrophage-mediated inflammation and an involvement of cross-presenting dendritic cells in the maintenance of an antiviral cytotoxic T cell response and virus control in the chronic infection phase. Together our data demonstrate a delicate adaptation process towards a chronic virus infection with both immunosuppressive and immunostimulatory processes. We fill a knowledge gap regarding the mechanisms of effector T cell maintenance and provide a new rational for targeted therapeutic vaccination.

## Introduction

Overwhelming infections with non-cytopathic viruses can lead to chronic infections. The fate of a virus infection can be fundamentally categorized as acute or chronic according to its temporal relationship with the host organism (1). In humans, acute infections are usually resolved within a few weeks. By contrast, chronic infections are not resolved and, instead, develop when innate and adaptive immune responses are not sufficient to eliminate the invading virus during the primary infection phase. Examples for this later case are infections with the Human Immunodeficiency Virus (HIV) or the Hepatitis B and C viruses (HBV, HCV) that can establish persistence in their hosts with different probabilities and pathogenic consequences (2,3).

A hallmark of an overwhelming infection is the downregulation of immune effector mechanisms to avoid immunopathology. Indeed, the simultaneous presence of a widespread virus infection and strong cytotoxic effector cell responses can induce massive cell and tissue destruction, and may directly threaten the life of the infected host (4). This threat is sensed by not completely understood mechanisms and several suppressive immune components are activated that adapt the host response to the viral threat. Amongst these are T cell exhaustion by deletion (5) and functional impairment (6), the generation of monocyte-derived suppressor cells (MDSCs) (7,8), and regulatory cell subsets (9). They exert their function via distinct processes including interaction with inhibitory ligands, the production of soluble immunosuppressive factors like IL10 and Indoleamine 2,3-dioxygenase, and the delivery of suppressive signals by cell-cell contacts to conventional T cells and to antigen-presenting cells thus influencing effector functions directly or indirectly (6,10–12).

Despite the various suppressive mechanisms induced during a chronic virus infection, the effector T cell shut-down is only partial and some T cell functionality remains that restrains the expansion of a persisting virus (13–16). The processes however that mediate the transition towards a lower level response are not well understood. We therefore sought to analyze on a systems level infection-fate–specific gene signatures and adaptive processes of the host to an overwhelming virological threat. The infection of mice with lymphocytic choriomeningitis virus (LCMV) was used as a model system. It enables to reproducibly establish acute and chronic infections, and exhibits several fundamental features within the virus-immune system’s crosstalk that hold true for human infections as well (17). Our results show, first, that spleen-derived transcriptomes cluster according to different virus infection stages thus validating the use of whole organ signatures as a valuable, phenotype correlating data set. Second, gene coexpression network analysis of time-resolved transcriptomes reveals that only few genes are involved in infection fate-specific biological pathways. Third, in chronic infection, there is an early attenuation of macrophage-mediated inflammation that occurs prior to the onset of T cell exhaustion. And finally, in the time window when CD8^+^ T cell exhaustion appears, there is a recruitment of cross-presenting XCR1^+^ dendritic cells that contributes to the maintenance of an antiviral cytotoxic T cell response and participates in viral control in the chronic infection phase. Together our data provide a mechanistic understanding of the adaptation towards an overwhelming virus infection with both immunosuppressive and immunostimulatory processes that leave options for therapeutic interventions.

## Results

### Time-resolved spleen transcriptomes cluster according to LCMV infection stages

To define and characterize on a systems level virus infection-fate-specific gene signatures, C57BL/6 mice were infected with a low-dose (2×10^2^ PFU; acute infection) or a high-dose (2×10^6^ PFU; chronic infection) of LCMV strain Docile (LCMV_Doc_), and analyzed longitudinally for virus titers, virus-specific CD4^+^ and CD8^+^ T cell responses and spleen-derived transcriptomes. Due to the different inoculum size, virus titers in spleen increased later in acute than in chronic infection, however similar levels were reached at d5 postinfection (p.i.) (Fig. 1A). As previously described (5), only low-dose infection resulted in virus clearance at d31 p.i.. Initial expansion of virus-specific IFN-ϒ-producing T cells showed a similar kinetics in acute and chronic infections, with a peak at d6-d7 p.i.. However, their number dropped at d7-d9 in high-dose infected mice which is indicative of CD8^+^ T cell exhaustion (Fig. 1B and Supplementary Fig. 1A). To characterize changes of transcriptional profiles from spleens during acute and chronic infections, we isolated total messenger RNAs (mRNAs) from naive mice (d0) and from mice at d3, d5, d6, d7, d9 and d31 after low-dose or high-dose of LCMV_Doc_ infection, and determined time-resolved transcriptomes by RNA-seq. Overall, the number of differentially expressed genes after acute and chronic infection reflected the virus dynamics (Supplementary Fig. 1B). Hierarchical clustering across all samples revealed four main groups corresponding to different infection phases (Fig. 1C). One group contains samples from acute infected mice at days 3 and 31, and uninfected mice (d0), indicating that spleen transcriptome profiles return to that of naive mice once the infection is resolved. Two groups represent early and late effector responses. Finally, d9 and d31 samples from chronic infection, that is once CD8^+^ T cell exhaustion is established, cluster in a separate group thus representing specifically the chronic infection phase. Together these data show that distinct virus infection phases exhibit separated transcriptome signatures in the lymphoid tissue.

**Fig 1.**
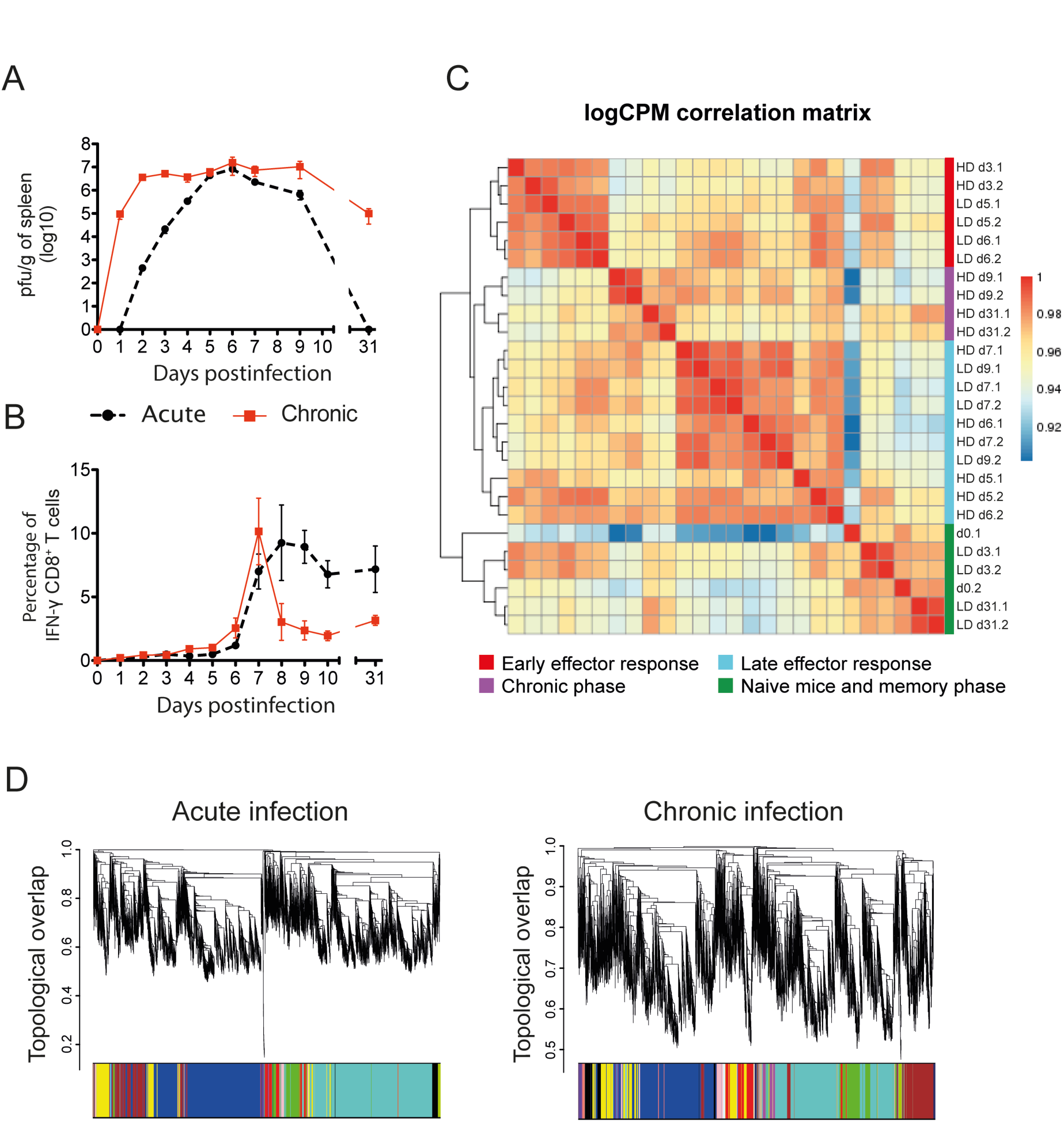
Kinetics of virus expansion and CD8^+^ T cell response, and transcriptome analysis in acute and chronic LCMV infections. (A and B) Virus titers (A) and percentages of GP33-specific IFN-ϒ-producing CD8^+^ T cells (B) in spleen. The mean ± SEM is shown. (C) Correlation matrix heatmap of gene expression across acute (LD) and chronic (HD) samples. The color scale indicates the degree of correlation. (D) Hierarchical clustering dendrogram for all differentially expressed genes (lines) obtained by WGCNA. The branches correspond to modules of highly coexpressed groups of genes. Colors below the dendrogram indicate the module that each gene was assigned to.

### Gene coexpression networks reveal distinguishing features of the adaptive immune responses in acute and chronic infection

To characterize the distinguishing traits between acute and chronic infections, genes of the time-resolved transcriptomes were grouped by weighted gene coexpression network analysis (WGCNA)(18). Topological overlap was used to measure the connection strength between genes based on shared network neighbours, identifying genes with similar patterns of connections and subsequently defining modules of highly coexpressed genes (Fig. 1D). Finally, the eigengene was extracted for each module to represent the gene expression profile. 23 color-coded modules were identified from acute and chronic infections (Supplementary Fig. 2), with a number of genes per module ranging from 15 to 4952 (Supplementary Fig. 3).

CD8^+^ T cells play a major role in the control of an LCMV infection as well as in exhaustion during chronic infections (19). Thus, to test if the obtained coexpression modules represent relevant biological pathways, we investigated whether the behavior of virus-specific T cells was mirrored by a gene expression pattern of similar kinetics. For this we correlated the module eigengenes with the kinetics of IFN-ϒ-producing GP33-specific CD8^+^ T cells (Fig. 2A). The acute-brown module significantly correlated with the virus-specific CD8^+^ and CD4^+^ T cell responses of acute infection (Fig. 2B). To visualize this correlation at the level of the individual genes, we compared the intramodular connectivity (K_IM_) of each gene of this module with its correlation to the T cell response. A correlation was observed indicating a link between the hub genes (high K_IM_ values) to the T cell response (Supplementary Fig. 4A). In contrast, no significant correlation between chronic-module eigengenes and the virus-specific CD8^+^ T cell response of chronic infection was observed (Fig. 2A). This indicates that IFN-ϒ production is a good marker to identify pathways involved in the T cell response in acute infection but not sufficient to reveal the underlying gene signature of exhaustion.

**Fig 2.**
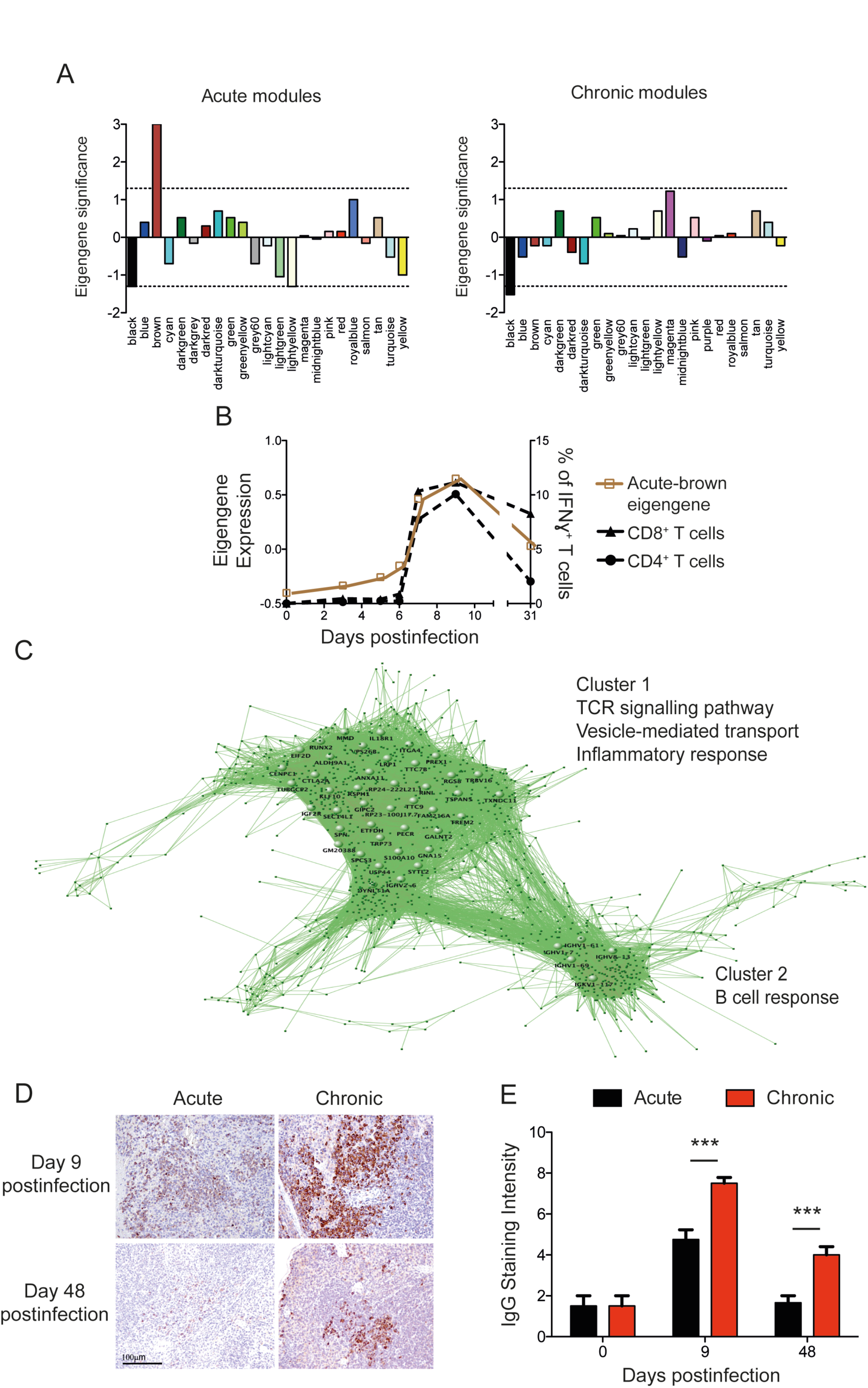
Genes related to T and B cell responses from the acute-brown module are disassociated in chronic infection. (A) Bar plot of eigengene significances indicating the correlation (Pearson) of each module with virus-specific IFN-ϒ-secreting CD8+ T cell kinetics in acute and chronic infection (-log(pvalue); modules with negative correlation are plotted with a negative value). (B) Acute-brown module eigengene and kinetics of GP33-specific CD8+ and GP61-specific CD4+ T cells. (C) Visualization of the acute-brown module genes. Large nodes represent hub genes with high connection densities (TOM>0.5). (D) Representative images of IgG immunohistochemical staining of spleens. Magnification bar: 100µm. (E) Semiquantification of IgG-positive cells in spleens (score 0-10). Data shown are mean ± SEM of 4 mice per group and time point. *** p ≤ 0.001 (unpaired two-tailed t test).

Hub genes from coexpression modules are defined by a high K_IM_ value, or alternatively by a high topological overlap, and are predicted to be central for the biological processes they represent (10,20). Thus, we visualized the hub genes of the acute-brown module as a network based on their topological overlap values (Fig. 2C). Two clusters were observed that contain genes involved in T cell activation (cluster 1) and B cell response (cluster 2). To analyse how these hub genes in acute infection behave during chronic infection, we compared their intramodular connectivities in both infection outcomes. Genes related to T cell activation were found in two different chronic modules, chronic-tan and chronic-greenyellow (Supplementary Fig. 4B). However, only genes in chronic-tan retained a K_IM_>0.6 and therefore represented control points of T cell activation in chronic infection. These genes were upregulated at day 6 and returned to basal levels at day 31 thus having a similar kinetics as in acute infection (Supplementary Fig. 4C). In contrast, genes in the chronic-greenyellow module showed an upregulation at day 6 with a subsequent downregulation at day 9 thus coinciding with T cell exhaustion. To validate these observations and to link the day 9 downregulation of the chronic-greenyellow genes to T cell exhaustion, we performed qPCR of the chronic-tan genes Cd247 and Cd3e, and the chronic-greenyellow genes Cd3d and ZAP70, that play a critical role in T cell receptor signaling, from spleens of mice at day 9 after acute or chronic infection (Supplementary Fig. 4E and F). The qPCR results positively correlated with the RNA-seq data. Cd3d and ZAP70 genes were significantly downregulated in chronic infection thus supporting a role during T cell exhaustion. The B cell response-related genes of cluster 2 were mainly found in the chronic-green module and retained a high intramodular connectivity value (Supplementary Fig. 4B). The expression kinetics of these genes during acute and chronic infections were identical, being upregulated from d7 to d9 and remaining elevated at d31. However, their expression levels were notably higher in chronic infection (Supplementary Fig. 4D and F) suggesting a causal link to the previously described hypergammaglobulinemia of chronic infections (21). To validate this, spleens from acute and chronic LCMV infections were stained for total IgG and analyzed by immunohistochemistry. IgG production was evident at d9 in both types of infection but was significantly higher in chronic infection. An elevated level of IgG remained visible even at day 48 postinfection when the B cell response of acute infection was comparable to that of naive mice (Fig. 2D and 2E). Taken together, coexpression network analysis from spleen-derived transcriptomes permits the identification of biologically relevant molecular pathways that are characteristic for acute or chronic infections.

### Early attenuation of the inflammatory response in chronic infection

To identify the common and the specific features for acute and chronic infection fates, we quantified module preservation (the degree to which coexpressed genes in acute infection are also coexpressed in chronic infection) by computing a connectivity-based preservation-statistic (Z-summary) that allows scoring the degree of similarity of the connectivity patterns between genes from two networks. Eigengene expression profiles of highly preserved modules between acute and chronic infections are shown in Supplementary Fig. 3B and C. Relevant overlapping hub genes between preserved modules are listed in Supplementary Table 1. Module preservation scores were inversely correlated with the percent of differentially connected genes (defined as those with a difference in their K_IM_ between acute and chronic networks higher than 0.4) (Supplementary Fig. 5A). This indicates that the preserved modules, that share similar gene connectivity patterns, also share common hub genes. Furthermore, module preservation scores correlated with the module size (Supplementary Fig. 5A), indicating that the biological pathways that are specific for an infection fate (low preservation scores) are governed by few genes. The kinetics of the identified five acute-specific and four chronic-specific modules are given in Supplementary Fig. 5B and C while their respective genes and connectivity values are listed in Supplementary Table 2. The acute-specific modules showed diverse expression profiles characterized by marked expression peaks at different time points. The included genes represent both innate and adaptive immune pathways highlighting the need of diverse biological responses to properly resolve a viral infection. On the other hand, all chronic infection-specific modules showed expression changes between days 7 and 9 p.i., when T cell exhaustion appears (Supplementary Fig. 5C), thus indicating that critical events governing chronic infection fates occur in this time window.

From the acute-specific modules, the acute-grey60 module was most interesting. It has a marked expression peak at day 6 postinfection just before the specific T cell response appeared thus bridging innate immune responses with adaptive responses. All 31 genes of this module are highly coexpressed only during acute infection (Fig. 3A) and enriched for genes involved in the regulation of IL6 and IL12 production (Supplementary Fig. 6A). As IL6 and IL12 are pro-inflammatory cytokines with important roles in antiviral host responses, our observations suggested a differential regulation in both infection fates before T cell exhaustion becomes apparent. To verify this hypothesis, we plotted the *Il6* and *Il12b* expression kinetics from our RNA-seq data (Supplementary Fig. 6B) and validated the IL6 expression results by qPCR including additional infected animals. Both, RNA-seq and qPCR gave consistent results showing a peak of IL6 expression at d6 after acute infection that was lacking in chronic infection (Supplementary Fig. 6C). Since inflammatory responses are complex responses, we next sought to identify whether *Il6* and *Il12* were part of a broader cluster of coexpressed genes within their acute-blue module. Indeed, we found 1504 genes in acute-blue module to correlate to the *Il6* expression kinetics with a correlation value above 0.8. Moreover, the level of expression of 79 of these genes was significantly higher in acute than in chronic infection at d6 p.i. (log2FC>1) (Fig. 3B). A list of 27 representative genes from this submodule, and their expression profiles in acute and chronic infections is shown in Supplementary Fig. 6D and in figure 3C, respectively. KEGG pathway analysis showed enrichment of genes involved in inflammatory pathways such as the NOD-like receptor and the Jak-STAT signaling pathways, and cytokine-cytokine receptor interaction. Moreover, we identified two genes related to the metabolism of Arginine and Proline (*Arg2* and *Nos2*), a pathway known to play an important role in the regulation of inflammation by monocytes and macrophages (22). *Il1b* and *Nos2* expression kinetics was further confirmed by qPCR including additional infected mice (Supplementary Fig. 6C). Together these data show differential spleen monocyte/macrophage characteristics during early time points of acute and chronic virus infections.

**Fig 3.**
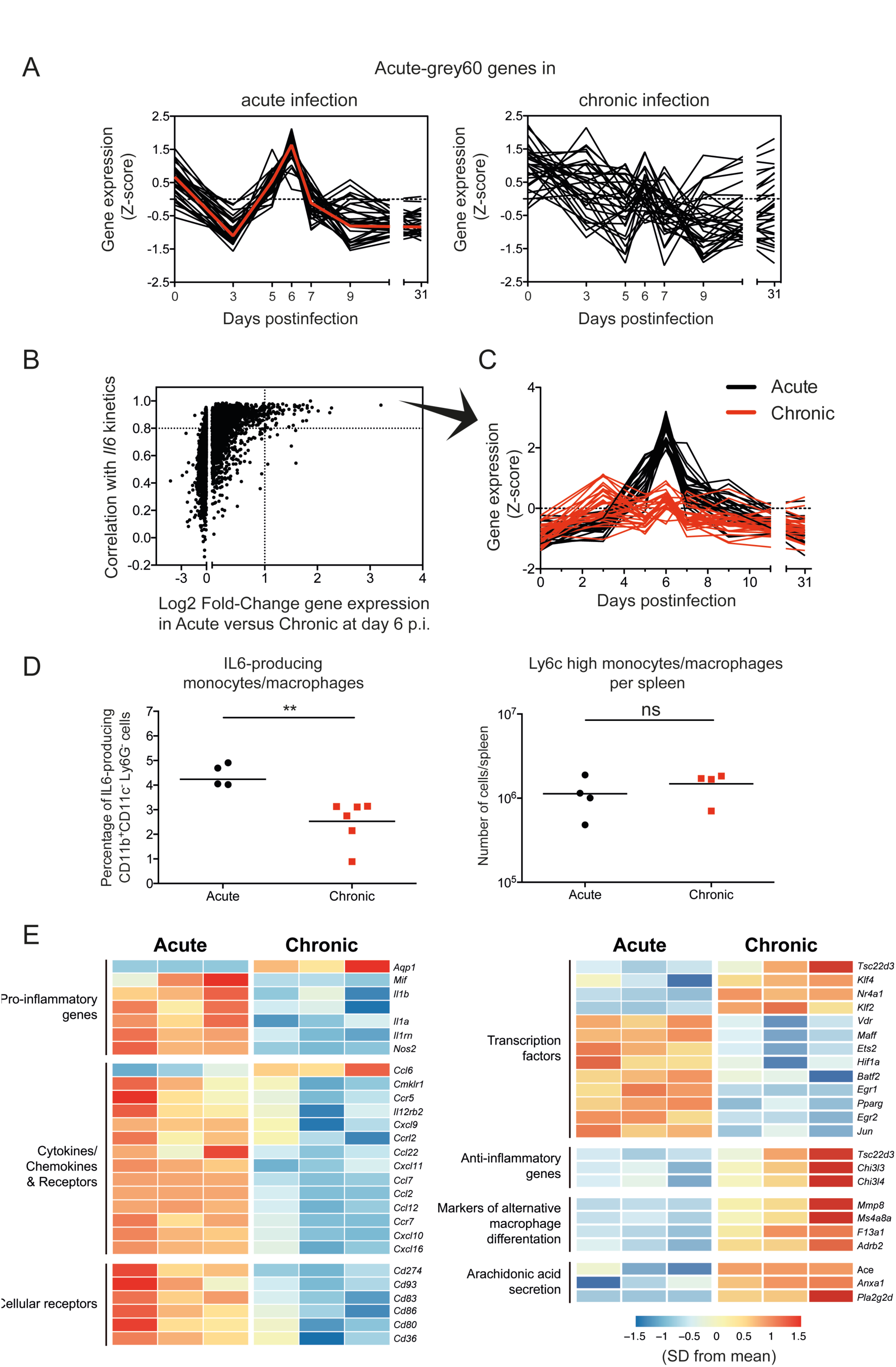
Early attenuation of inflammatory monocytes/macrophages in chronic infection. (A) Normalized expression kinetics of genes from the acute-grey60 module and their corresponding kinetics behaviour in acute infection. Red line represents the kinetics of the eigengene. (B) Pearson’s correlation between genes of the acute-blue module and *Il6* in acute infection is plotted against the log2 fold-change (log2FC) of gene expression between acute and chronic infection at day 6 p.i.. Dashed lines mark a correlation >0.8 and log2FC>1. (C) List of 27 representative genes with a correlation>0.8 with *Il6* expression kinetics, and log2FC>1 of gene expression between acute and chronic infection at day 6 p.i.. Grey boxes indicate the enriched KEGG pathways that each gene belongs to. (D) Percentage of IL6-producing CD11b^+^ CD11c^-^ Ly6G^-^ cells and number of Ly6C high monocytes/macrophages per spleen at day 6 postinfection. Data shown are mean ± SEM of n=4 to 5 mice. * p ≤ 0.05; ** p ≤ 0.01; *** p ≤ 0.001 (unpaired two-tailed t test). (E) Heatmap illustrating the relative expression of genes in monocyte/macrophage cells from acute and chronic infected mice at day 6 p.i..

To test the hypothesis of an early differential polarization of monocyte/macrophages during both outcomes of LCMV infection, we characterized spleen monocyte/macrophages at day 6 postinfection by intracellular IL6 staining and RNA-seq. The percentage of IL6-producing monocytes/macrophages from chronic infection were significantly lower than those from acute infection while the number of pro-inflammatory Ly6C^hi^ monocytes/macrophages in the spleen were, as described before (7), similar in both infections (Fig. 3D). Transcriptional profiling from sorted monocytes/macrophages showed that cells from acute infection expressed higher levels of pro-inflammatory cytokines and chemokines, and cellular receptors and transcription factors associated to M1 macrophages. In contrast, monocytes/macrophages from chronic infection expressed higher levels of transcription factors and genes that exert anti-inflammatory effects and markers of M2 macrophage differentiation (Fig. 3E). These data demonstrate that inflammatory monocytes/macrophages are induced early after an acute LCMV infection and, in contrast, during chronic infection these cells shift to an anti-inflammatory profile before T cell exhaustion becomes evident.

### The XCL1-XCR1 communication axis links T cell exhaustion with effector cell maintenance

Cytokines and chemokines are major components of the signalling network that regulates the coordinated immune response against a viral infection (23). Thus, we next investigated which cytokines/chemokines-encoding hub genes were present in acute- and chronic-specific modules, and therefore might participate in processes involved in infection fate regulation. From all the identified hub genes (Supplementary Table 2), of particular interest was the chemokine-encoding gene *Xcl1* from the chronic-darkturquoise module. XCL1 is mainly produced by NK and activated CD8^+^ T cells, and promotes the recruitment of dendritic cells expressing the receptor XCR1 (XCR1^+^ DCs) which are implicated in the priming and boosting of cytotoxic responses to cross-presented antigens (24,25).

All 17 genes of the chronic-darkturquoise module are highly coexpressed in chronic infection and deregulated in acute infection (Fig. 4A). They show a “two-peak” behaviour with an expression peak at d5 and a second upregulation from d7 to d9 p.i.. The second peak of expression of XCL1 was verified by qPCR using additional infected mice (Supplementary Fig. 7A). In contrast to chronic infection, *Xcl1* showed only a single early peak of expression at d6 in acute infection, returning to basal levels from d7 p.i. (Fig. 4B). To investigate the role of XCL1 during the establishment of the chronic infection phase, we first analyzed its expression by NK and CD8^+^ T cells at d9 by intracellular staining. Both cell types showed a higher expression of XCL1 in chronic infection compared to acute infection or uninfected mice, however CD8^+^ T cells were the main producers (Fig. 4C and Supplementary Fig. 7B). Importantly, XCL1 was highly produced by LCMV-specific CD8^+^ T cells once exhaustion was established (Fig. 4D), and particularly abundant in the subpopulation of CXCR5^+^ cells (Fig. 4E). It has recently been shown that virus-specific CXCR5^+^ CD8^+^ T cells play a major role in the control of virus replication during chronic infections (13,14,26). Our data now show that they also play an important role in the immune adaptation process during exhaustion.

**Fig 4.**
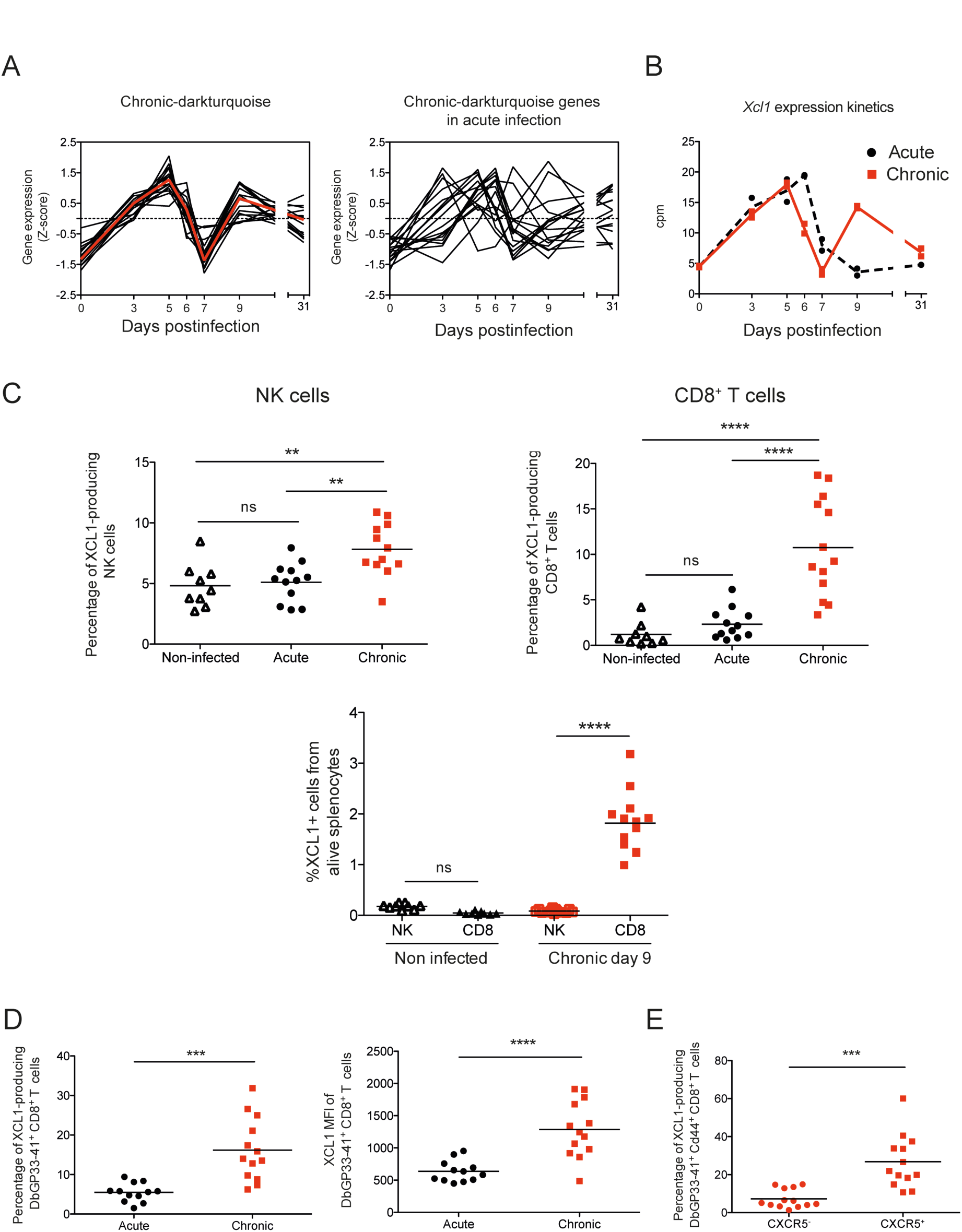
Chronic-darkturquoise specific module reveals an important role of XCL1 in chronic infection. (A) Normalized expression kinetics of genes from the chronic-darkturquoise module and their corresponding kinetics behaviour in acute infection. (B) *Xcl1* expression kinetics obtained from the RNA-seq analysis. (C) Percentages of XLC1-producing NK and CD8^+^ T cells at day 9 p.i. Data shown are mean ± SEM from n= 11 to 13 mice representative of three independent experiments (significance determined using one-way ANOVA). (D and E) MFI of XLC1 in DbGP33-41^+^ CD8^+^ T cells (D) and percentages of XLC1-producing DbGP33-41^+^ CD8^+^ T cells (D) and CXCR5^-^ or CXCR5^+^ DbGP33-41^+^ CD44^+^ CD8^+^ T cells (E) at day 9 p.i.. Data shown are mean ± SEM from n= 11 to 13 mice (significance determined using unpaired two-tailed t test). ns, not significant; ** p ≤ 0.01; *** p ≤ 0.001; **** p ≤ 0.0001.

Coinciding with the second peak of XCL1 expression, the quantification of XCR1^+^ DCs showed a clear increase of these cells in the spleens at d9 in chronic infection (Fig. 5A and Supplementary Fig. 7C). To further analyze the impact of this XCL1-XCR1 axis, we depleted XCR1^+^ DC in chronically-infected XCR1-DTRvenus mice (27) with diphtheria toxin (DT) (Fig. 5B and Supplementary Fig. 7D). While the virus-specific CD4^+^ T cell response was not affected, XCR1^+^ DC depletion led to a significant reduction in the percentage of GP33-specific CD8^+^ T cells at day 15 p.i. and lower percentages of CD107a, CD107b and IFN-ϒ producing cells than in untreated animals (Fig. 5C). Moreover, the percentage of CXCR5^+^ CD8^+^ T cells was also significantly decreased, as well as the percentages of CD107a/b producing CXCR5^+^ CD8^+^ T cells (Fig. 5D), demonstrating an interdependence between this CD8^+^ T cell subset and cross-presenting XCR1^+^ DCs. Consistently, the viral titers in spleen, lung and kidney were higher in mice lacking XCR1^+^ DCs (Fig. 5E). Together these results demonstrate that the XCL1-XCR1 communication axis is (i) an important component of immune system adaptation to an overwhelming virus threat and (ii) an important component of virus control in the chronic infection phase.

**Fig 5.**
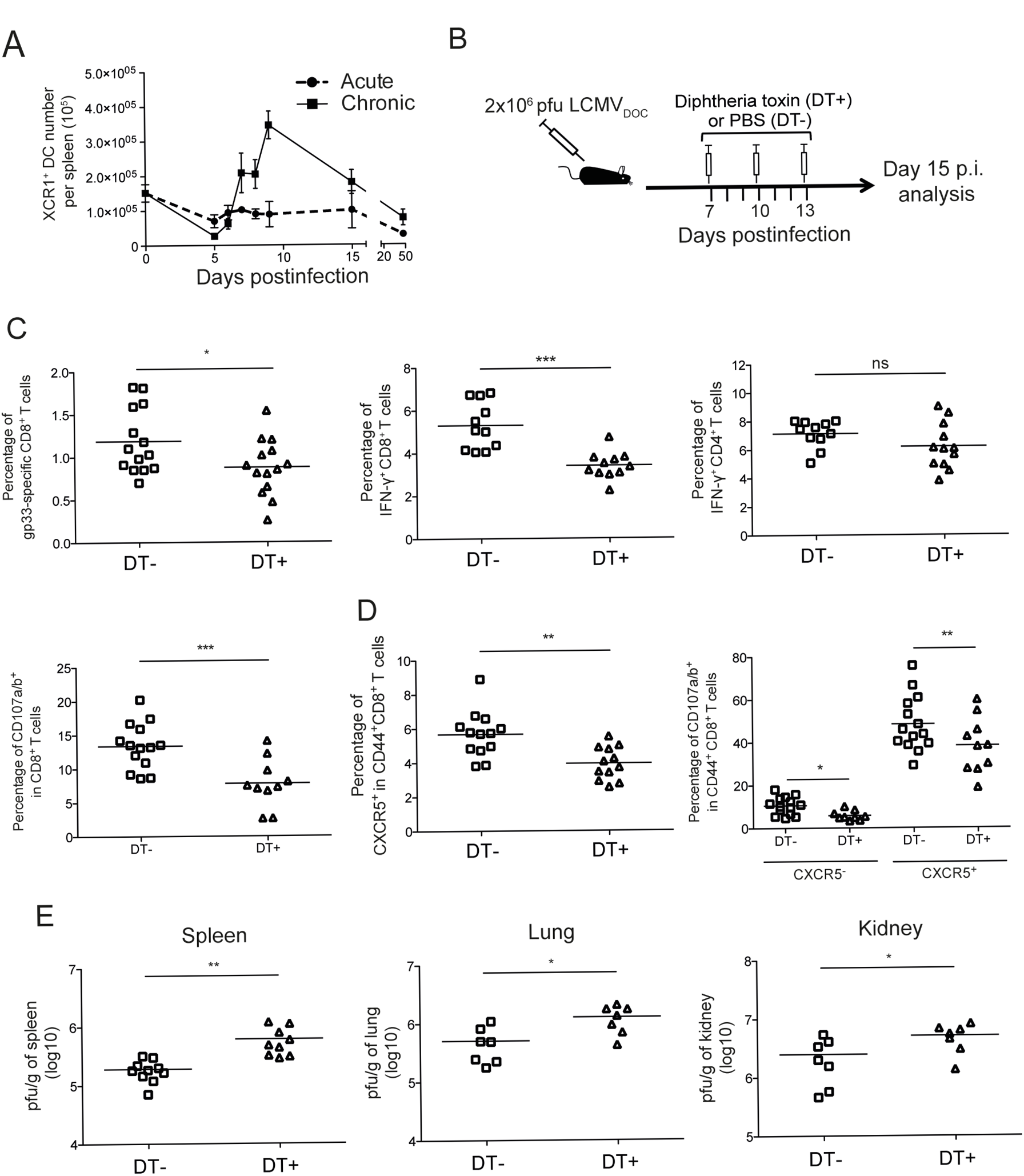
XCR1^+^ dendritic cells are required to maintain antiviral T cell response and control virus titers. (A) Number of XCR1^+^ dendritic cells in spleen. (B) Schematic representation of DT treatment regimen for depletion of XCR1^+^ dendritic cells. C Percentages of DbGP33-41^+^ CD8^+^ T cells, IFN-ϒ-producing CD8^+^ and CD4^+^ T cells, CD107a^+^ and CD107b^+^ CD8 T cells, and CXCR5^+^ CD44^+^ CD8^+^ T cells from chronic infected mice non treated (DT−) or treated (DT+) with DT at day 15 p.i.. (D) Virus titers in spleen, lung and kidney from chronic infected mice non treated (DT−) or treated (DT+) with DT at day 15 p.i.. (C and D) For each group replicates and the mean ± SEM is shown. ns, non significant; * p ≤ 0.05; ** p ≤ 0.001; *** p ≤ 0.0001 (unpaired two-tailed t test). Data are representative of two or three independent experiments.

### Discussion

Here we used time-resolved spleen transcriptomes from acutely or chronically-LCMV-infected mice in combination with weighted gene coexpression network analysis to decipher differentiating host responses and biological pathways. We demonstrate an early attenuation of inflammatory monocyte/macrophage responses prior to T cell exhaustion and the involvement of the XCL1-XCR1 communication axis in virus containment during chronic infection (Fig. 6). Thus, the response of a host to an overwhelming infection threat is complex and subtle, involving both immune-stimulatory and immune-suppressive pathways.

**Fig 6.**
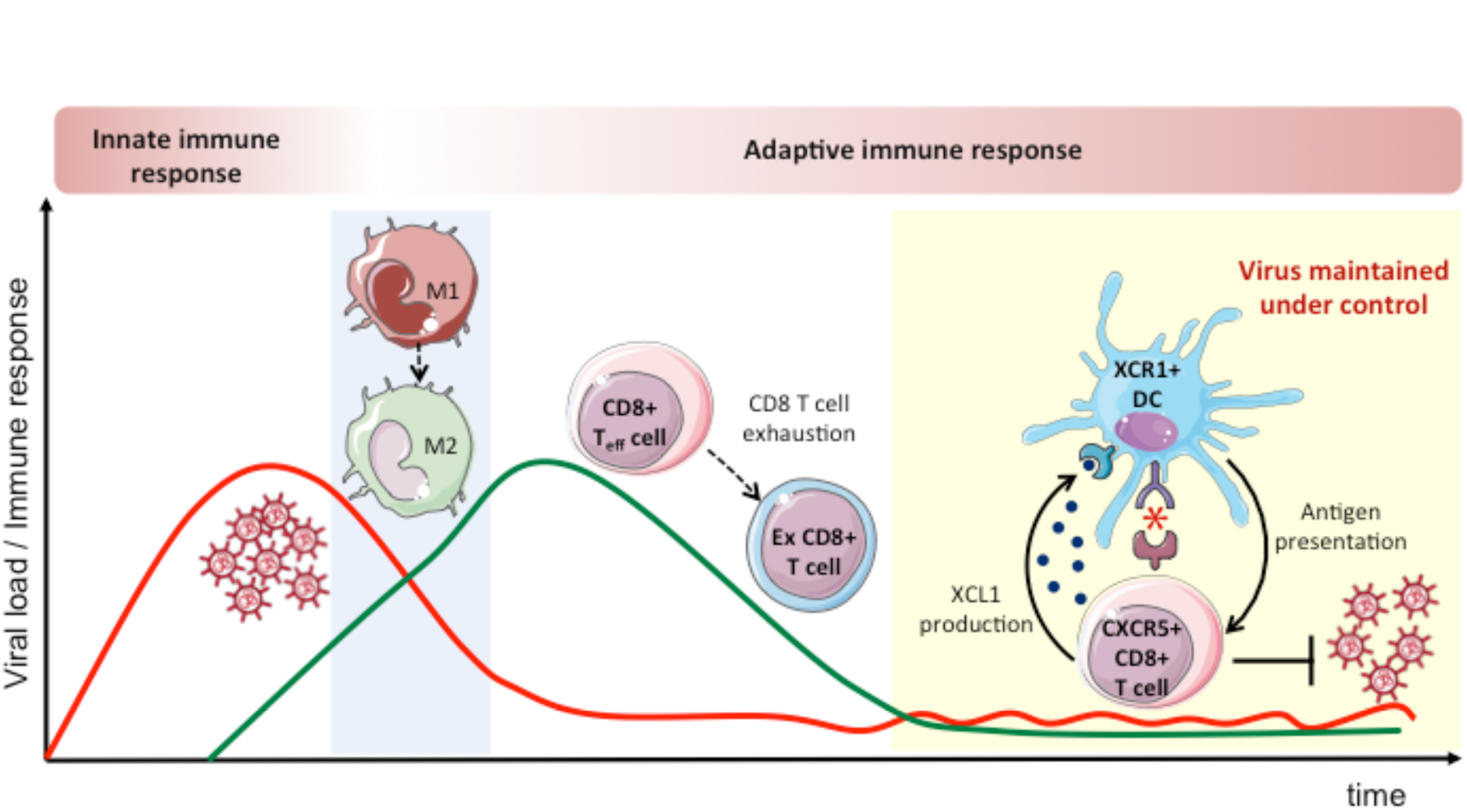
Schematic representation of the host adaptation towards a chronic virus infection. The early inflammatory macrophage attenuation and the role of the XCL1-XCR1 communication axis in virus containment during the chronic infection phase is shown. Images were adopted from Servier Medical Art by Servier (https://smart.servier.com/) and modified by the authors under the following terms: CREATIVE COMMONS Attribution 3.0 Unported (CC BY 3.0)(https://creativecommons.org/licenses/by/3.0/).

Acute and chronic infection fates are readily distinguished by their sequential splenic transcriptomes. The whole transcriptomes reveal differentiating traits at day 9 postinfection when acute infection starts to be resolved. However, close inspection of the coexpression modules demonstrates an attenuation of inflammatory monocytic cells at day 6 before T cell exhaustion becomes apparent. Thus, the decision point to downregulate immune responses towards an overwhelming virus threat is already taken before T cells shut off their effector functions. Whether this decision point lies even earlier remains to be determined. So far, early innate immune responses represented i.e. by modules containing type I IFN genes are very similar in acute and chronic infections, and differences come up only from day 6 onwards. In addition to these global characteristics, we were also able to link specific genes and pathways to critical, already established phenotypes associated with both infection fates including the effective LCMV-specific T cell response as well as T cell exhaustion and hypergammaglobulinemia. Together this validates our systems approach for analyzing virus infection fates and leaves a rich comprehensive data set for further, more specific analyses.

Infection-fate-specific module information provides a firm basis to reveal specific, physiologically relevant changes in the infected host organism. An intriguing observation was that the infection-fate-specific modules contained rather few genes. This notion may not only help to define diagnostic biomarker for predicting infection outcomes but also hint to regulators of pathways that may direct phenotype evolution over time. Indeed, the acute-grey60 module with its 31 genes that have an expression peak at day 6, is a good example. Starting from a GO enrichment analysis of its genes we subsequently identified important differences in early monocytic cell polarization with an M1-type profile in acute infection and cells with an attenuated phenotype during chronic infection. Previous studies have described a skewing of monocytic cells towards MDSCs in chronic LCMV infections of mice (7) and in HIV, HBV and HCV infections in humans (8,28,29). MDSCs appear during the primary infection phase and have been characterized, i.e. by transcriptome analysis at day 14 post-chronic LCMV clone 13 infection, a few days after T cell exhaustion becomes apparent (7). Our own RNA-seq results from isolated monocytic cells at day 6 postinfection now point to an even earlier phenotypic shift with an expression of genes linked to alternative macrophage differentiation and anti-inflammation.

Concomitant with CD8^+^ T cell exhaustion during chronic infection, XCR1^+^ DC number increase in spleen. Depletion of XCR1^+^ DCs resulted in reduction of virus-specific CD8^+^ T cells and a concurrent increase in viral loads (Fig 5). However, as downregulation of T cell effector function by exhaustion is a mechanism to avoid immunopathology (30), as is attenuation of inflammatory macrophages and generation of MDSCs (31), why should a host organism at the same time expand a subtype of antigen-presenting cells that are highly potent in priming and boosting novel T cell responses? This fundamental issue may simply highlight two important aspects within the *in vivo* virus-immune system cross-talk. First, the immune system has to shut down when confronted with an overwhelming viral threat. A high-dose LCMV_Doc_ infection requires an adaptation of the immune response and viral load dynamics such that it levels below a life-threatening situation. This is achieved by a variety of immune suppressive mechanisms of which T cell exhaustion and the generation of MDSCs are important components. Second, the immune system does not surrender completely to an overwhelming threat but maintains partial control and restricts virus overloads. One possible way for this seems to be a feedback regulation involving cells from the population of virus-specific CD8^+^ T cells that become exhausted. Indeed, the population of LCMV-specific CD8^+^ T cells that becomes exhausted contains cells that produce XCL1 (Fig. 5D) and thus can set up XCR1^+^ DC recruitment and cytotoxic T cell stimulation and effector cell maintenance. Overall, this argues for the operation of virus-threat-sensing host mechanisms that adapt the immune system effectors to minimize damage and the XCL1-XCR1 axis of being part of a functional adaptation to the chronic infection phase.

How may these observations be exploited for host benefit? Recent publications have shown that virus-specific CXCR5^+^ CD8^+^ T cells expand most efficiently after immune checkpoint inhibition (13,14). They are, besides NK cells, the main producer of XCL1 in the chronic infection phase (Fig. 4C). Furthermore, XCL1 is elevated in HIV-infected elite controllers

(32) and its receptor XCR1 is solely expressed on cross-presenting DCs (33) that are specifically upregulated during the chronic infection phase (Fig. 5A and Supplementary Fig. 7C). Therefore, enforcing the XCL1-XCR1 axis in an immunotherapeutic intervention seems a promising strategy to better control chronic virus infections. One option in this direction would be the already established targeting of immunogens to XCR1^+^ DCs via linkage to XCL1 itself or to a XCR1-targeting antibody (34,35). Combining such immunotherapeutic vaccination with checkpoint inhibition may then allow effector cell expansion enhanced by a positive feedback loop and thus a greater increase in the effector cell to virus ratio.

In conclusion, coexpression network analysis of time-resolved splenic transcriptome data revealed fundamental features of infection fate regulation. Several links between gene transcripts and immunological phenotypes were established that include an early attenuation of inflammatory monocytic cells and an involvement of the XCL1-XCR1 axis in the functional adaptation of the immune response in a chronic virus infection. This not only provides a unique data set to generate and test novel hypotheses about immune system functioning but also suggests directions for immunotherapeutic strategies to better control or cure persistent infections. An inspection of the results in human infections and cancer is clearly indicated.

## Materials and Methods

### Animals, infections, and *in vivo* cell depletion

Six- to twelve-week-old C57BL/6 (male) (RRID:IMSR_JAX:000664) and XCR1-DTRvenus mice (male and female) (27) were infected intraperitoneally (i.p.) with either a low dose (LD; 2×102 plaque forming units) or a high dose (HD; 2×106 plaque forming units) of the strain Docile of LCMV (LCMV_Doc_) to induce an acute or chronic infection, respectively. Viral titers from spleens of infected mice were determined on MC57 cells using focus-forming assay (36). For depletion of XCR1+ dendritic cells, XCR1-DTRvenus mice (heterozygote and control mice) were infected with a high dose of LCMV to induce a chronic infection, and Diphtheria Toxin (DT; 25ng/g body weight) was administered at days 7, 10 and 13 postinfection. Physiological serum was used as control vehicle. Mice were purchased from Charles River Laboratories and maintained under specific-pathogen-free conditions at in-house facilities.

#### Ethics statement

All experimental procedures utilizing mice were conducted at the animal facility of the Barcelona Biomedical research Park (PRBB) according to the guidelines from Generalitat de Catalunya (project number 9422) and approved by the ethical committee for animal experimentation at Parc de Recerca Biomèdica de Barcelona (CEEA-PRBB, Spain; permit license number JMR-16-0046). Mice were euthanized by carbon dioxide inhalation.

### Flow cytometry, sorting, and immunohistochemistry

For Flow Cytometry analysis and sorting of cells, spleens were harvested and single-cell suspensions were generated. Biotinylated MHC class I monomer (MBL International Corporation) was tetramerized using PhycoPro R-PE conjugated to Streptavidin (Prozyme). For determination of IL6-producing monocytes/macrophages, splenocytes from half of each spleen were directly put into media containing Brefeldin A without stimulation before Intracellular Cytokine Staining (ICS). All antibodies were purchased from either BD Biosciences, eBioscience or Biolegend. Data were analyzed using FlowJo 10.1 software. A LSR Fortessa (BD Biosciences) was used for flow cytometry. Monocytes/macrophages and DbGP33-41^+^ CD8^+^ T cells were sorted in a FACSAria II SORP (BD Biosciences). All samples were kept at 4ºC during the sorting and sort purity was >95% for all populations. Spleen samples for immunohistochemistry were embedded in paraffin after an overnight fixation with 4% buffered formaldehyde. Three micrometer thick tissue sections were deparaffinized and rehydrated before staining with Mayer’s Hematoxylin and Eosin. Extra sections were obtained and were incubated with DAKO EnVision+ System HRP labelled polymer with anti-Mouse IgG for immunohistochemical labelling and semiquantification of IgG-positive cells. The reaction was visualized using hydrogen peroxide and 3-3’-diaminobenzidine as a chromogen substrate. A set of scores was defined from 0 (absence of immunolabeled cells in the studied sections) to 10 (100% IgG labelled cells in the studied sections). For semiquantification of fibrosis, spleen sections were stained with Masson’s Trichrome staining, which stains collagen fibers in color blue. A set of values was defined from 0 to 10: a score of 0 represents normality (a certain amount of blue stained collagen is to be observed in the splenic capsule and trabeculae); a score of 10 would correspond to a spleen in which the parenchyma has totally been replaced by connective tissue.

### RNA-seq library preparation and sequencing

Total RNA from spleens (15-20 mg) and sorted cells (5×104 cells per sample) was isolated according to the manufacturer’s instructions using Qiagen RNeasy Mini kit and Qiagen RNeasy Micro kit (Qiagen), respectively. RNA was submitted for sequencing to the Genomics Unit of Centre for Genomic Regulation (CGR, PRBB). The quality and concentration of RNA were determined by an Agilent Bioanalyzer. Sequencing libraries were obtained after removing ribosomal RNA by a Ribo-Zero kit (Illumina). cDNA was synthesized and tagged by addition of barcoded Truseq adapters. Libraries were quantified using the KAPA Library Quantification Kit (KapaBiosystems) prior to amplification with Illumina’s cBot. Four libraries were pooled and sequenced (single strand, 50nts) on an Illumina HiSeq2000 sequencer to obtain 50-60 million reads per sample.

### RNA-seq bioinformatic analysis

Reads mapping against the Mus musculus reference genome (GRCm38) was done using the GEMtools RNA-seq pipeline (http://gemtools.github.io/docs/rna_pipeline.html), and were quantified with Flux Capacitor (http://sammeth.net/confluence/display/FLUX/Home) with the Mus musculus gencode annotation M2 version (https://www.gencodegenes.org/). Normalization was performed with the edgeR TMM method(37). Pairwise Pearson’s correlation coefficients (PCC) were calculated for comparison among transcriptomes of spleens from uninfected (n=2, day 0), acute (n=2 per time point) or chronic (n=2 per time point) infected mice. Hierarchical clustering across all samples was based on pairwise Pearson’s correlation coefficients among RNA-seq libraries. Differential expression analysis was performed with the ‘robust’ version of the edgeR R package(38). Genes with a false discovery rate (FDR)<5% were considered significant. Differentially expressed genes in acute and chronic time series (n=13971) were used to construct two signed coexpression networks with the WGCNA R package for each dataset (39). First, a signed weighted adjacency matrix was calculated with the ‘blockwiseModules’ function using these parameters: power=30, TOMtype="signed", minModuleSize=15, mergeCutHeight=0.25, reassignThreshold=0, networkType="signed", numericLabels=TRUE, pamRespectsDendro=FALSE, nThreads=7, maxBlockSize=17000. The power law of 30 was selected to meet the scale-free topology assumption with the pickSoftThreshold function. Then, genes were clustered into network modules using average linkage hierarchical clustering and the topological overlap measure (TOM) as proximity. Each of the identified modules was summarized by its module eigengene (the first principal component), which represents the weighted average expression profile of all module genes (39). To identify intramodular hub genes inside a given module, the intramodular connectivity was calculated for each gene (Kin) and ViSANT (http://visant.bu.edu/) was employed for network visualizations (TOM>0.3). Module preservation and module overlapping were calculated with functions ‘modulePreservation’ and‘userListEnrichment’, respectively. Viral loads, CD4 and CD8 levels were correlated with the module eigengenes with Pearson correlation. Gene ontology (GO) enrichment analysis was performed with DAVID (http://david.ncifcrf.gov/)(40).

### Module preservation analysis

Z-summary, implemented within WGCNA, with 100 random permutations of the data, is used as a connectivity-based preservation statistic able to determine whether the connectivity pattern between genes in a reference network is similar to that in a test network. A Z-summary score < 2 indicates no evidence of preservation, 2 < Z-summary < 10 implies weak preservation and Z-summary > 10 suggests strong preservation. Highly preserved modules were defined as those with a preservation Z-summary value above 10 and percentage of differentially connected genes below 20%. Group-specific modules were defined as those with a preservation Z-summary value below 2 and a percentage of differentially connected genes above 35%.

### Quantitative Real-Time PCR

Total RNA from spleens was used to perform quantitative real-time PCR using SYBR select master mix (ThermoFisher). cDNA templates were generated using SuperScript III reverse transcriptase (ThermoFisher). Quantitative real-time PCR was performed in a Quantstudio 12K flex (ThermoFisher). Expression of all target genes was normalized to expression of the housekeeping gene Gapdh. Primers for all genes were designed with Primer Express 3.0 (Applied Biosystems). Primer sequences are as follows: Gapdh forward, 5’-CCA GTA TGA CTC CAC TCA CG-3’ and reverse, 5’-GAC TCC ACG ACA TAC TCA GC-3’; Xcl1 forward, 5’-TTT GTC ACC AAA CGA GGA CTA AA-3’ and reverse, 5’-CCA GTC AGG GTT ATC GCT GTG-3’; Il6 forward, 5’-TGG GAC TGA TGC TGG TGA CA-3’ and reverse, 5’-TTT CCA CGA TTT CCC AGA GAA-3’; Il1b forward, 5’-GGA GCT CCC TTT TCG TGA ATG-3’ and reverse, 5’-TCT TGG CCG AGG ACT AAG GA-3’, Nos2 forward, 5’-ACT CTT CAC CAC AAG GCC ACA T-3’ and reverse, 5’-GTT GAT GAA CTC AAT GGC ATG AG-3’; ZAP70 forward, 5’-TCG GCA CTA TGC CAA GAT CA-3’, and reverse, 5’-TCA CTG CGG CTG GAG AAC TT-3’; CD3d forward. 5’-GTG GAA GGA TGG TTT GCA AA-3’ and reverse, 5’-CAC ACA GTT CTG GCA CAT TCG-3’; CD3e forward, 5’-CCA GCG GGA CCT GTA TTC TG-3’, and reverse, 5’ - AAC AAG GAG TAG CAG GGT GC-3’; Fut8 forward, 5’-GTT ATT GGA GTC CAT GTC AG-3’, and reverse, 5’-TTG GAG TAC TTT GTC TTT GC-3’; Ighg2c forward, 5’-TGA TTG TGC AGA CCC TCG TG-3’, and reverse, 5’-CTG TGG ACT GGA CCA GCA AT-3’; Cd247 forward, 5’-GCT GGA TCC CAA ACT CTG CT-3’, and reverse, 5’-GCT GTT TGC CTC CCA TCT CT-3’.

### Statistical analysis

Two-tailed t test or one-way ANOVA analyses were performed using GraphPad Prism 6.0 (San Diego, CA, USA). pvalues (p) below 0.05 were considered significant and were indicated by asterisks: *p ≤ 0.05; **p ≤ 0.01; ***p ≤ 0.001; ****p ≤ 0.0001. Non-significant differences were indicated as “ns”. Fisher’s exact test was used to quantify overlap between modules.

### Data Availability

The complete RNA-seq datasets are available from the Gene Expression Omnibus (accession number GSE107880).

## Supporting information

## Acknowledgments

We like to thank Mónica Pérez and Rita De Giuli-Buehrer for their excellent technical support.

