## Supplementary material for "Time-resolved systems analysis reveals a critical role of XCR1+ dendritic cells in the maintenance of effector T cells during chronic viral infection"

**S1 Table. Overlapping hub genes between preserved modules.**

When the number of overlapping genes>100, only genes representing relevant biological functions obtained from David are shown.

| Acute Yellow | # Overlapping genes: 159 | Chronic Brown |
|---|---|---|

Innate immune response

*Oas3, Bst2, Mx2, Herc6, Zbp1, Ddx58, Oas1a, Ifit2, Pml, Ifih1, Ripk2, Mx1, Ifitm3, Samhd1, Oasl1, Rnf135, Ifit3, Irf7, Trim21, Dhx58, Ifit1*

Cellular response to interferon-alpha

*I830012O16Rik, Ifit3, Myc, Ifit2, Ifit1*

Cellular response to interferon-beta

*Trex1, Ifit3, Ifi205, Ifit1*

Antigen processing and presentation of peptide antigen via MHC class I

*H2-T22, H2-Q4, H2-M3, H2-T23, H2-Q6, H2-Q7, H2-K1, H2-T10*

| Acute Turquoise | # Overlapping genes: 108 | Chronic Yellow |
|---|---|---|

Regulation of transcription, DNA-templated

*Rfx7, Zfp62, Purb, Suv420h1, Zfp81, Zscan26, Ubp1, Zfp68, Pura, Zfp560, Zfp317, Zfp770, Txlng, Rnf38, Gm12258, Ehf, Pou2f1*

Positive regulation of GTPase activity

*Gm13304, Rapgef6, Dennd6a, Ccl21a, Gm21541*

Lymphocyte chemotaxis

*Gm13304, Ccl21a, Gm21541*

| Acute Turquoise | # Overlapping genes: 541 | Chronic Blue |
|---|---|---|

Regulation of transcription, DNA-templated

*Crtc1, Jarid2, Zfp764, Zscan2, Sap25, Brd3, Kdm5b, Mapk11, Zfp318, Asxl3, Zfp688, Zfp579, Il16, Gm9897, Zfp932, Zfp943, Ring1, Arid3b, Brwd1, Wasl, Zbtb9, Zfp866, Gpbp1l1, Zfp740, Foxp1, Krba1, Zbtb49, Ebf4, Hes5, Phf1, Smad7, Rfx1, Prdm11, Akap8l, Zfp420, Cux1, Zfp653, AW146154, Zfp867, Crebzf, Ebf1, Srsf5, Sp6, Cbx7, Gm15446, Mterfd2, Crtc3, Hmg20a, Zfp942, Rarg, Foxj3, Kdm3b, Epc1, Cdkn2aip, Hes6, Zfp212, Ezh1, Ccnt2, Kdm4b, Chd6, E4f1, Ccar2, Hif1an, Arid4b, Camta1, 5830428H23Rik, Foxj2, Nfe2l3, Zfp512, Ddx5, Per3, Meis1, Phf21a, Sirt7, Zfp182, Dbp, Zfp592, Tle2, Zmym5, Dmtf1, Mterf, Nfrkb, Zfp275, Zfp746, Zscan20, Fam120b, Irf3, Hdac10, Akap8, Tspyl2, Zfp956, Prdm9, Lrif1, Mterfd3*

Covalent chromatin modification

*Jarid2, Phf21a, Hmg20a, Brd3, Ing4, Sirt7, Kdm5b, Phf1, Kdm3b, Epc1, Ring1, Ogt, Hdac10, Ezh1, Tspyl2, Chd6, Kdm4b, Msl1, Cbx7, Prdm9*

Protein phosphorylation

*Map2k7, Sbk1, Obscn, Ulk3, Stk11, Phkg2, Map3k1, Mapk11, Ikbkb, Clk4, Prkcg, Map4k2, Clk1, Mast3, Nek8, Tec, Pan3, Clk2, Map4k1, Ltk, Fgfr3, Npr2, Csnk1g2*

| Acute Blue | # Overlapping genes: 669 | Chronic Turquoise |
|---|---|---|

Cell cycle

*Tipin, Ran, Uhrf1, Clspn, Kif2c, Zwilch, Mastl, Gm6531, Ska3, Suv39h1, Ckap2, Cks2, Gmnn, Mybl2, Cdc20, Aurka, Cenpw, Cdc34, Cdca8, Nuf2, Cdk4, Psrc1, E2f7, Mcm8, Ercc6l, Pin1, Cinp, Rbbp8, Ncaph, Mcm4, Ccnb1, Mcm6, Chaf1a, Erh, Cdc123, Cdc25a, Bub1b, Mad2l1, Gsg2, Cep55, Nasp, Ube2s, Spdl1, Cdca2, Cdk1, Spc24, Dbf4, Nup37, Tfdp1, Tpx2, Lig1, Cdca5, Plk1, Cks1b, Mcm2, Bub1, Aurkb, Ppm1g, Hells, Ticrr, Chaf1b, Mcm5, Mcm3, Cdc45, Melk, Nek2, Bub3*

<table>
<tr><th>Acute Blue</th><th># Overlapping genes: 36</th><th>Chronic Magenta</th></tr>
<tr><td colspan="3">Itgb2, Lcp2, Por, Kctd5, Fos, Slc7a8, Adamts1, Slc16a3, Ccr1, Slc11a1, Tagln2, Soat1, Rbms1, Plin2, Ptpn12, Itgam, F7, Pkm, Pdcd6ip, Alas1, Atp1a1, Rnpep, Hilpda, D630023F18Rik, C5ar1, Rnd1, Gm15590, Aldoart1, Atp8b4, Gm12164, Gm13365, Mrgpra2a, Gm13812, Gm13810, Mrgpra2b, E230013L22Rik</td></tr>
<tr><th>Acute Green</th><th># Overlapping genes: 16</th><th>Chronic Yellow</th></tr>
<tr><td colspan="3">Unc5b, Cacna1g, Kat6b, Slc12a2, Dcaf12, Gm996, 2410066E13Rik, Zfhx3, Pitpnc1, Foxo4, A830010M20Rik, B230219D22Rik, Mmp12, Sox4, Zfp433, Gm26702</td></tr>
<tr><th>Acute Brown</th><th># Overlapping genes: 16</th><th>Chronic Tan</th></tr>
<tr><td colspan="3">Cd247, Grb7, Fam71b, Prkch, Tmem180, Tubgcp2, Cd3e, Gipc2, Kremen2, Themis, Spn, Trbv14, Trbv29, Zan, B930095G15Rik, 4930519F09Rik</td></tr>
<tr><th>Acute Green</th><th># Overlapping genes: 11</th><th>Chronic Pink</th></tr>
<tr><td colspan="3">Trpv4, Abcc3, Acot2, Fam168a, Mgll, Tie1, Decr2, Snrk, Fndc4, Fam211b, Tmem19</td></tr>
</table>

<table>
<tr><th>Acute Red</th><th># Overlapping genes: 13</th><th>Chronic Salmon</th></tr>
<tr><td colspan="3">Dip2a, Abca2, Tnik, 2610020H08Rik, Xlr4c, Galnt6, Cep250, Yeats2, Kif21b, Usp48, Lsm11, Xlr4b, Tmppe</td></tr>
</table>
