## Supplementary material for "Time-resolved systems analysis reveals a critical role of XCR1+ dendritic cells in the maintenance of effector T cells during chronic viral infection"

**S2 Table. Chronic-specific module gene list.**

| Acute_Magenta | |
|---|---|
| Gene | $K_{IM}$ |
| *Adgb* | 1 |
| *Serpina3b* | 0.978 |
| *Car12* | 0.947 |
| *Stk35* | 0.933 |
| *Myo19* | 0.924 |
| *Fdxr* | 0.911 |
| *Slc19a1* | 0.894 |
| *Lonp1* | 0.869 |
| *Dusp1* | 0.847 |
| *Ccrn4l* | 0.821 |
| *A930007I19Rik* | 0.813 |
| *Thbs1* | 0.808 |
| *Bcl2l1* | 0.802 |
| *Bend4* | 0.801 |
| *Doc2a* | 0.795 |
| *Pfas* | 0.778 |
| *Gm11508* | 0.758 |
| *Jmjd6* | 0.756 |
| *Plk2* | 0.722 |
| *Rassf1* | 0.71 |
| *Gm13199* | 0.705 |
| *Rbm14* | 0.696 |
| *Ampd2* | 0.68 |
| *Mrps10* | 0.671 |
| *Htr1b* | 0.668 |
| *Mettl13* | 0.656 |
| *Oxnad1* | 0.653 |
| *Mogs* | 0.637 |
| *Gm16229* | 0.617 |
| *Lrdd* | 0.606 |
| *Smpd4* | 0.602 |
| *Tcirg1* | 0.6 |
| *1700030J22Rik* | 0.599 |
| *Neu1* | 0.598 |
| *Cstf2* | 0.598 |
| *Furin* | 0.59 |
| *Slc7a6* | 0.585 |
| *Selp* | 0.585 |
| *Gm12201* | 0.571 |
| *Mrpl37* | 0.564 |
| *Pusl1* | 0.561 |
| *2410004B18Rik* | 0.553 |
| *Nab2* | 0.527 |
| *Armc6* | 0.508 |
| *Kbtbd8* | 0.494 |
| *Ddx11* | 0.491 |
| *Gm13904* | 0.483 |
| *Exoc1* | 0.479 |
| *Csf2ra* | 0.479 |
| *Zfp948* | 0.463 |
| *Gm16194* | 0.462 |
| *Tbrg4* | 0.452 |
| *Phlda1* | 0.449 |
| *Tsfm* | 0.443 |
| *Rab44* | 0.427 |
| *Adam12* | 0.424 |

| Acute_Magenta (cont) | |
|---|---|
| Gene | $K_{IM}$ |
| *Pkd1l3* | 0.412 |
| *1700034H15Rik* | 0.403 |
| *Wdr19* | 0.388 |
| *Grtp1* | 0.358 |
| *Cebpg* | 0.355 |
| *Kcnf1* | 0.337 |
| *3110043O21Rik* | 0.326 |
| *Tra2a* | 0.31 |
| *Papd5* | 0.289 |
| *Mafg* | 0.272 |
| *Sh3bp1* | 0.271 |
| *Scamp3* | 0.241 |
| *Epha2* | 0.223 |
| *Plekhm1* | 0.22 |
| *Lhfpl2* | 0.126 |
| *Lad1* | 0.077 |

| Acute_Darkturquoise | |
|---|---|
| Gene | $K_{IM}$ |
| *Fut4* | 1 |
| *Rp2h* | 0.963 |
| *Itm2c* | 0.923 |
| *Vcam1* | 0.893 |
| *Gm26924* | 0.87 |
| *Gm12913* | 0.865 |
| *Nxt2* | 0.842 |
| *Gm15564* | 0.695 |
| *Prcp* | 0.668 |
| *Ccl5* | 0.654 |
| *Rit1* | 0.649 |
| *Tpgs1* | 0.626 |
| *B3galnt1* | 0.588 |
| *Igkv1-122* | 0.481 |
| *Rab18* | 0.471 |
| *Pld3* | 0.454 |
| *Pparg* | 0.357 |
| *Gm8902* | 0.33 |
| *10-sept* | 0.244 |

| Acute_Darkgrey | |
|---|---|
| Gene | $K_{IM}$ |
| *Gm9200* | 1 |
| *2010204K13Rik* | 0.998 |
| *Rplp1-ps1* | 0.989 |
| *Gm26377* | 0.944 |
| *Psma8* | 0.883 |
| *Habp2* | 0.834 |
| *Uqcrh* | 0.82 |
| *Gm6206* | 0.792 |
| *1810043H04Rik* | 0.689 |
| *Rpp14* | 0.627 |
| *Siglece* | 0.569 |
| *Fis1* | 0.522 |
| *Gm11307* | 0.491 |
| *Emc6* | 0.353 |
| *Lyrm4* | 0.124 |

| Acute_Grey60 | |
|---|---|
| Gene | $K_{IM}$ |
| *Nufip2* | 1 |
| *Lilra6* | 0.93 |
| *Slc39a2* | 0.87 |
| *Camsap1* | 0.859 |
| *Traf4* | 0.784 |
| *Hgs* | 0.734 |
| *Nt5dc3* | 0.726 |
| *Cdk12* | 0.68 |
| *Trak1* | 0.666 |
| *Usp40* | 0.659 |
| *Hspa1l* | 0.61 |
| *Sfmbt2* | 0.587 |
| *Ier2* | 0.558 |
| *Lin37* | 0.549 |
| *Cass4* | 0.522 |
| *Slc29a2* | 0.521 |
| *Slc5a3* | 0.501 |
| *Arrdc4* | 0.501 |
| *Phf8* | 0.471 |
| *Pim3* | 0.442 |
| *Gm15228* | 0.434 |
| *Cpeb4* | 0.43 |
| *Urb1* | 0.398 |
| *Irak3* | 0.398 |
| *Usp3* | 0.387 |
| *Tlr6* | 0.384 |
| *Zfp655* | 0.366 |
| *Crlf2* | 0.338 |
| *Smox* | 0.329 |
| *Plekhm2* | 0.228 |
| *Gm16175* | 0.112 |

| Chronic_Lightgreen | |
|---|---|
| Gene | $K_{IM}$ |
| *Gm11747* | 1 |
| *Drwd1* | 0.982 |
| *C1qtnf1* | 0.857 |
| *Fam83g* | 0.819 |
| *Tmem9* | 0.81 |
| *Nudt9* | 0.805 |
| *Serp1* | 0.784 |
| *Capn5* | 0.74 |
| *Unc50* | 0.72 |
| *Fem1a* | 0.688 |
| *Oaf* | 0.653 |
| *Fstl1* | 0.639 |
| *Krt14* | 0.637 |
| *Mrgprf* | 0.613 |
| *B3gnt9* | 0.604 |
| *Psmf1* | 0.602 |
| *Kdelr3* | 0.58 |
| *Selk* | 0.563 |
| *Msln* | 0.556 |
| *Asf1a* | 0.555 |
| *Pla2g4c* | 0.507 |
| *Ahrr* | 0.504 |
| *Col18a1* | 0.493 |
| *Gm16174* | 0.485 |
| *Upk1b* | 0.439 |
| *Atp9a* | 0.435 |
| *Gm12475* | 0.396 |
| *Lax1* | 0.355 |
| *Zfp449* | 0.34 |
| *Mgat1* | 0.275 |
| *Cd248* | 0.261 |
| *Prune2* | 0.201 |
| *Mocs1* | 0.193 |
| *Plbd2* | 0.193 |
| *Tusc1* | 0.143 |
| *Dnajb1* | 0.101 |

| Chronic_Darkred | |
|---|---|
| Gene | $K_{IM}$ |
| *Plagl1* | 1 |
| *Slc9b2* | 0.942 |
| *Slamf6* | 0.905 |
| *Cx3cl1* | 0.897 |
| *Lrba* | 0.856 |
| *Idi1* | 0.831 |
| *Slain1* | 0.699 |
| *Sqle* | 0.64 |
| *Tnfsf11* | 0.623 |
| *Gm13502* | 0.607 |
| *Paip2b* | 0.6 |
| *Ap1ar* | 0.578 |
| *Drc1* | 0.559 |
| *Cyp51* | 0.553 |
| *Lss* | 0.525 |
| *Trbv12-1* | 0.491 |
| *Etv5* | 0.465 |
| *Ccdc141* | 0.451 |
| *Snap47* | 0.324 |
| *Igfbp2* | 0.321 |
| *Angptl7* | 0.311 |
| *Smco4* | 0.296 |
| *Fam26e* | 0.184 |
| *Fgf7* | 0.124 |
| *Adcy1* | 0.104 |

| Chronic_Darkturquoise | |
|---|---|
| Gene | $K_{IM}$ |
| *Gpr176* | 1 |
| *Xcl1* | 0.966 |
| *Metazoa_SRP* | 0.96 |
| *Metazoa_SRP* | 0.956 |
| *Kirrel* | 0.932 |
| *BC048507* | 0.875 |
| *Gm24105* | 0.802 |
| *Rpph1* | 0.78 |
| *Rai14* | 0.643 |
| *Gm22179* | 0.618 |
| *Taf5l* | 0.606 |
| *Enah* | 0.395 |
| *Dnase1l3* | 0.19 |
| *1700007L15Rik* | 0.158 |
| *Rpl5-ps1* | 0.131 |
| *Eif3s6-ps1* | 0.13 |
| *Trbv12-2* | 0.079 |

| Chronic_Darkgreen | |
|---|---|
| Gene | $K_{IM}$ |
| *Esr1* | 1 |
| *Gstm4* | 0.999 |
| *Fam188b* | 0.949 |
| *Lrrn1* | 0.922 |
| *Rit1* | 0.807 |
| *Galnt18* | 0.657 |
| *AB124611* | 0.635 |
| *Dcxr* | 0.622 |
| *Pafah1b1-ps1* | 0.621 |
| *Mgst1* | 0.545 |
| *Tmem50a* | 0.535 |
| *Bin3* | 0.493 |
| *H2-T3* | 0.442 |
| *Ppp1r2* | 0.387 |
| *Hpcal1* | 0.352 |
| *Pnpla2* | 0.317 |
| *1700009P17Rik* | 0.242 |
| *H2afv* | 0.237 |
| *Vkorc1* | 0.083 |
| *Esr1* | 1 |

| Acute_Darkgreen | |
|---|---|
| Gene | $K_{IM}$ |
| *Mfap3* | 1 |
| *4930524O07Rik* | 0.996 |
| *Gm16001* | 0.976 |
| *Arsg* | 0.911 |
| *Miat* | 0.875 |
| *Gm24187* | 0.875 |
| *Gm23935* | 0.873 |
| *Gm24245* | 0.873 |
| *Gm24270* | 0.867 |
| *Hoxa5* | 0.805 |
| *Zzef1* | 0.773 |
| *Pld2* | 0.76 |
| *Plek2* | 0.591 |
| *Etohi1* | 0.533 |
| *Hoxa3* | 0.498 |
| *Clec4b2* | 0.486 |
| *Gm13031* | 0.481 |
| *Hoga1* | 0.267 |
| *Zfp518b* | 0.263 |
| *Trio* | 0.177 |
