## Supplementary material for "Time-resolved systems analysis reveals a critical role of XCR1+ dendritic cells in the maintenance of effector T cells during chronic viral infection"

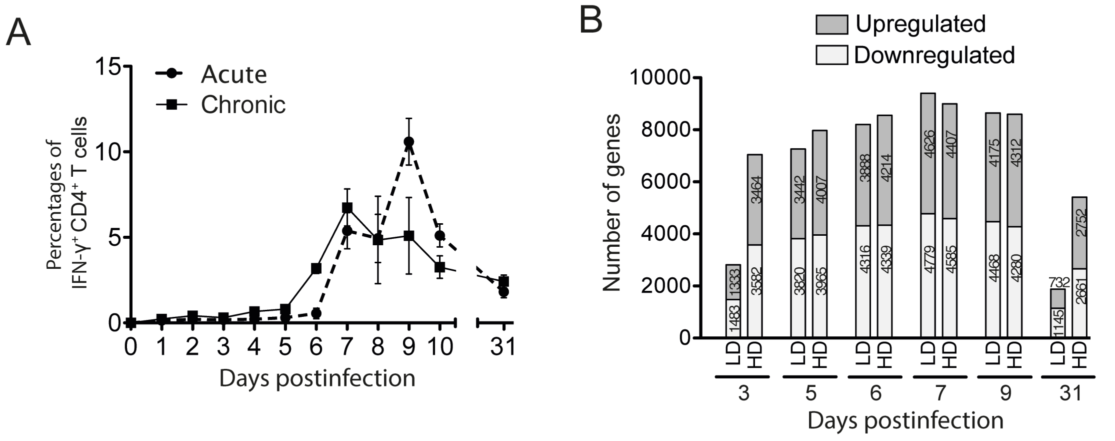


**S1 Figure. Kinetics of CD4+ T cell response and analysis of differentially expressed genes.** (A) Percentages of IFN-ϒ-producing $CD4^+$ T cells were determined by ICS after stimulation with GP61 peptide. For each group and time point, the mean ± SEM (standard error of the mean) is shown. (B) Differentially expressed genes in spleens after acute or chronic LCMV infections. Number of differentially expressed genes that were up- or down-regulated after infection were calculated for each time point.

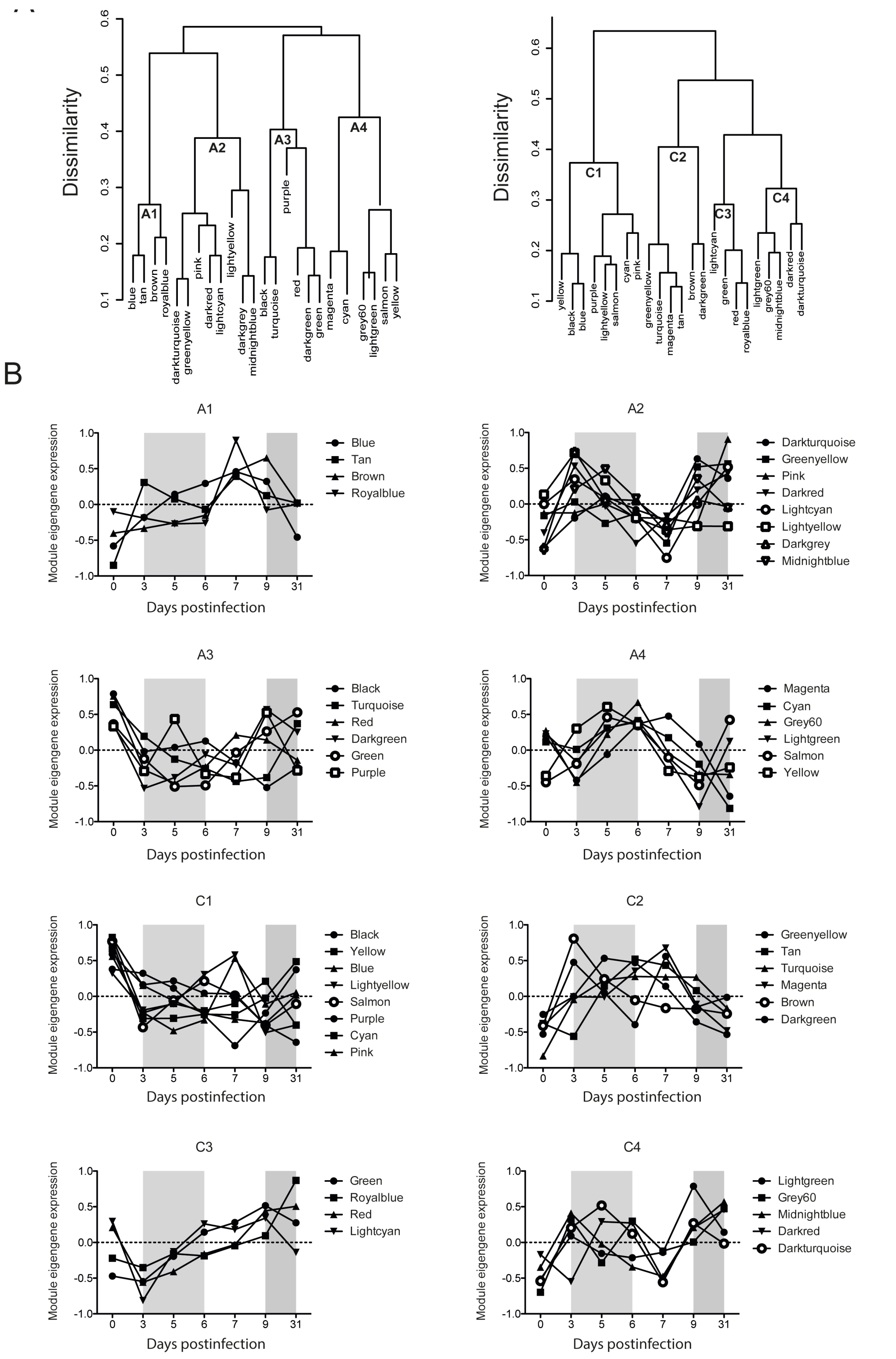


**S2 Figure. WGCNA-derived module eigengene expression patterns.** (A) Hierarchical clustering dendrogram of dissimilarity based on module eigengenes. To note, the same colour in modules from acute and chronic networks does not imply any similarity between them. (B) Module eigengene kinetics grouped based on hierarchical clustering obtained from acute (A) and chronic (C) infections.

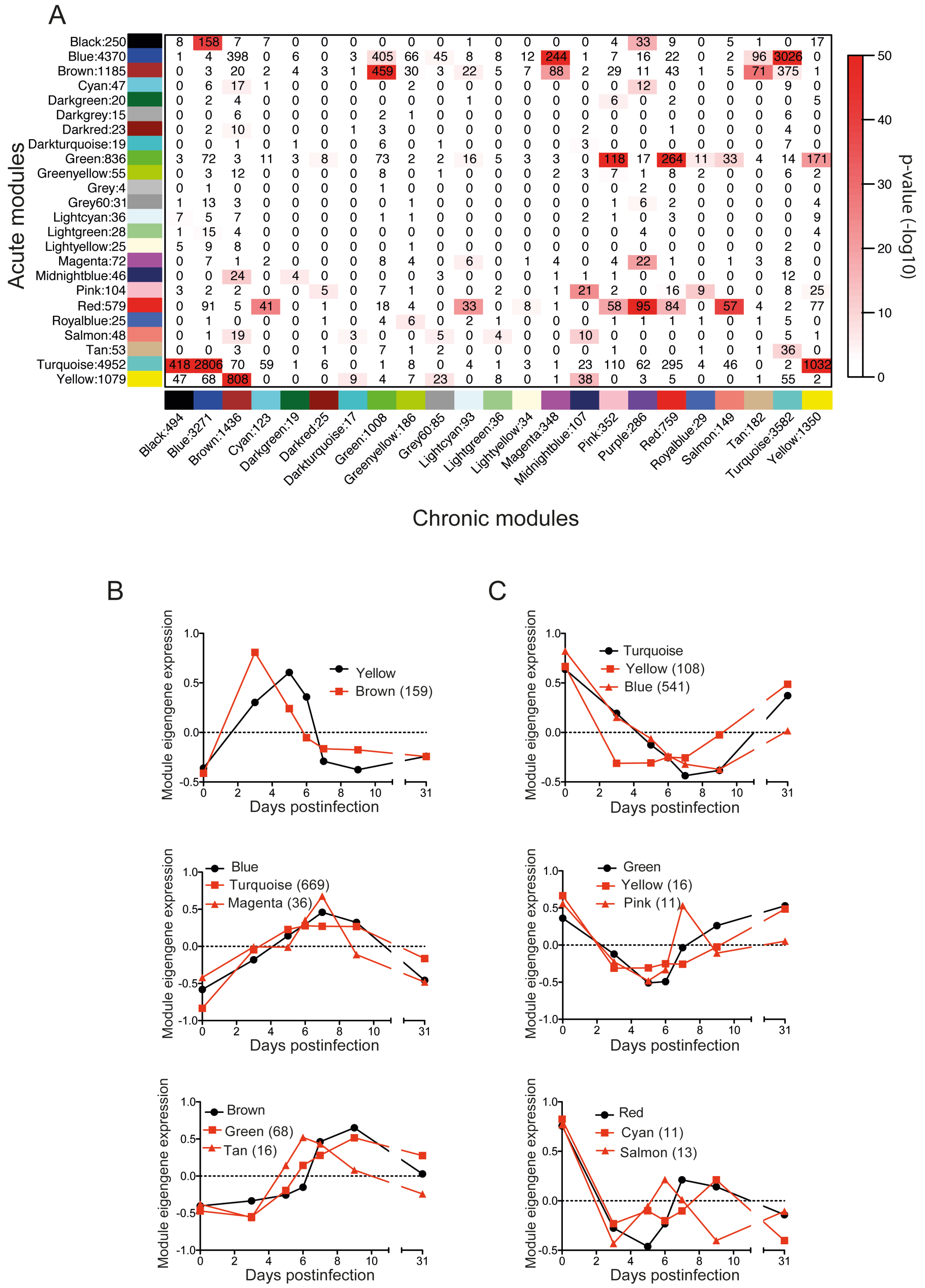


**S3 Figure. Acute and chronic module gene overlapping and expression kinetics of highly preserved modules.** (A) Heatmap showing module gene overlapping between acute and chronic infections. Each row and column is labeled by the corresponding module color, and the total number of genes in the module. Numbers within the table represents the number of genes in common between the corresponding acute and chronic modules. The significance of gene overlap was calculated by Fisher's exact test. Pvalues are color coded according to the color bar on the right. (B-C) Eigengene expression profiles of highly preserved modules between acute (red) and chronic (black) infections containing genes upregulated (B) or downregulated (C) after infection are shown. The number of overlapping hub genes between the corresponding chronic and acute modules are indicated in brackets.

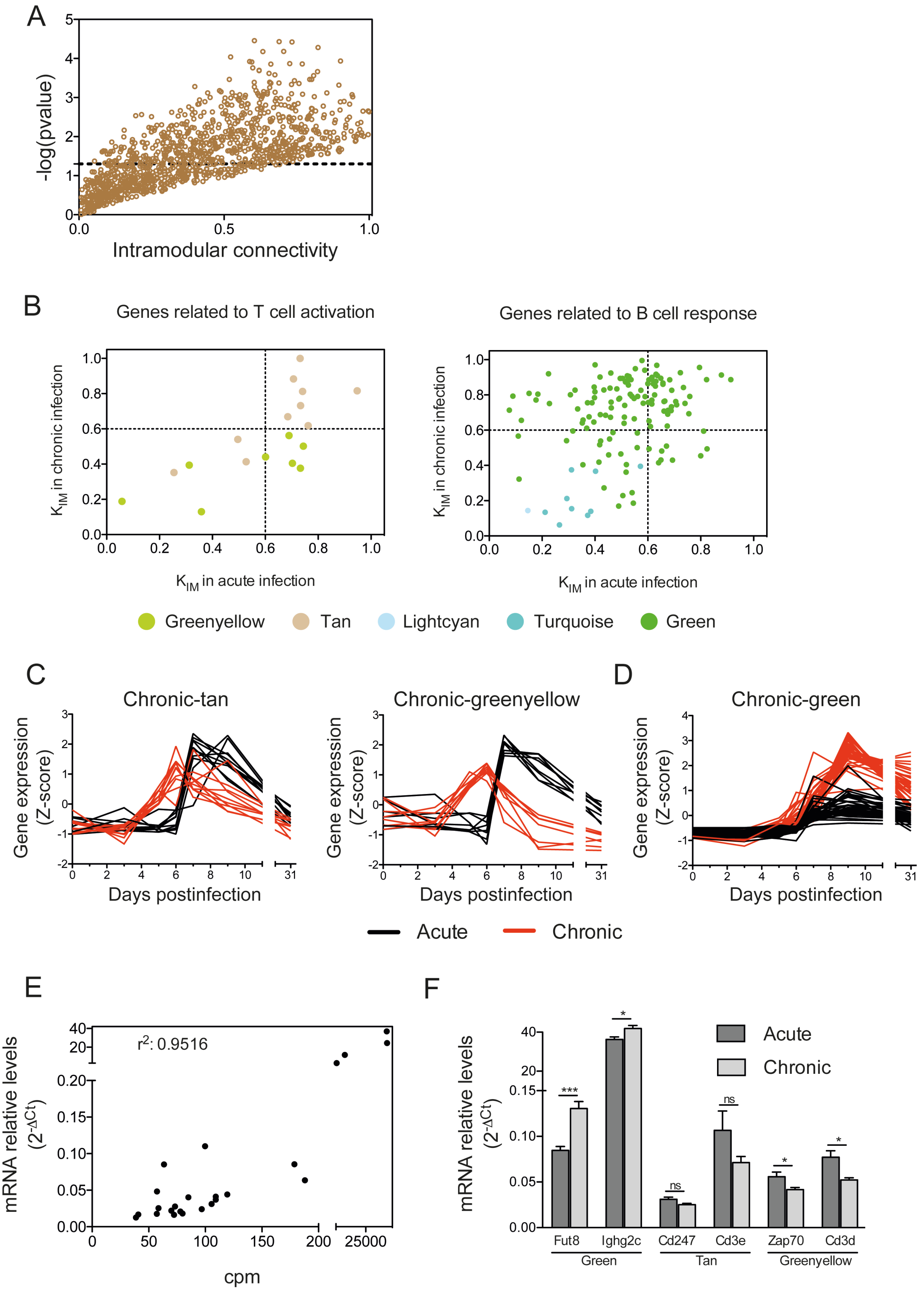


**S4 Figure. Acute-brown module represents T and B cell responses in acute infection.** (A) Intramodular connectivity versus GP33-specific CD8+ T cell kinetics correlation significance for acute-brown module genes. (B) Intramodular connectivity ($K_{IM}$) of genes related to T and B cell responses from the acute-brown module in acute versus chronic infection. Each gene is labeled by the corresponding module color of the chronic infection. Genes related to T cell activation are *Spn*, *Cd247*, *Cd3e*, *Cd3d*, *Themis*, *Prkcq*, *Zap70*, *Trbc1*, *Thy1*, *Grb7*, and the transcription factors *Maf* and *Nfatc2*; representative genes related to B cell response are *Fut8*, *Ighv*, *Iglv*, *Igkv*, *Igkc*, *Ighg*, *Ighj*, *Cdkn2c*, and Cdk6. (C-D) Expression kinetics of acute-brown module genes related to T cell activation (C) and hub genes related to the B cell response (D) in acute and chronic infections. (E-F) Expression levels at d9 p.i. of six genes related to T and B cell responses selected from acute-brown module were measured by qPCR and correlated to the expression values obtained by RNAseq in the same animals (E), and validated in a higher number of animals (F; 6 to 8 mice/group; in X-axis are shown gene names and their respective module in chronic infection). Significant differences were determined by an unpaired two-tailed t test. ns, non significant; * $p \leq 0.05$; *** $p \leq 0.001$.

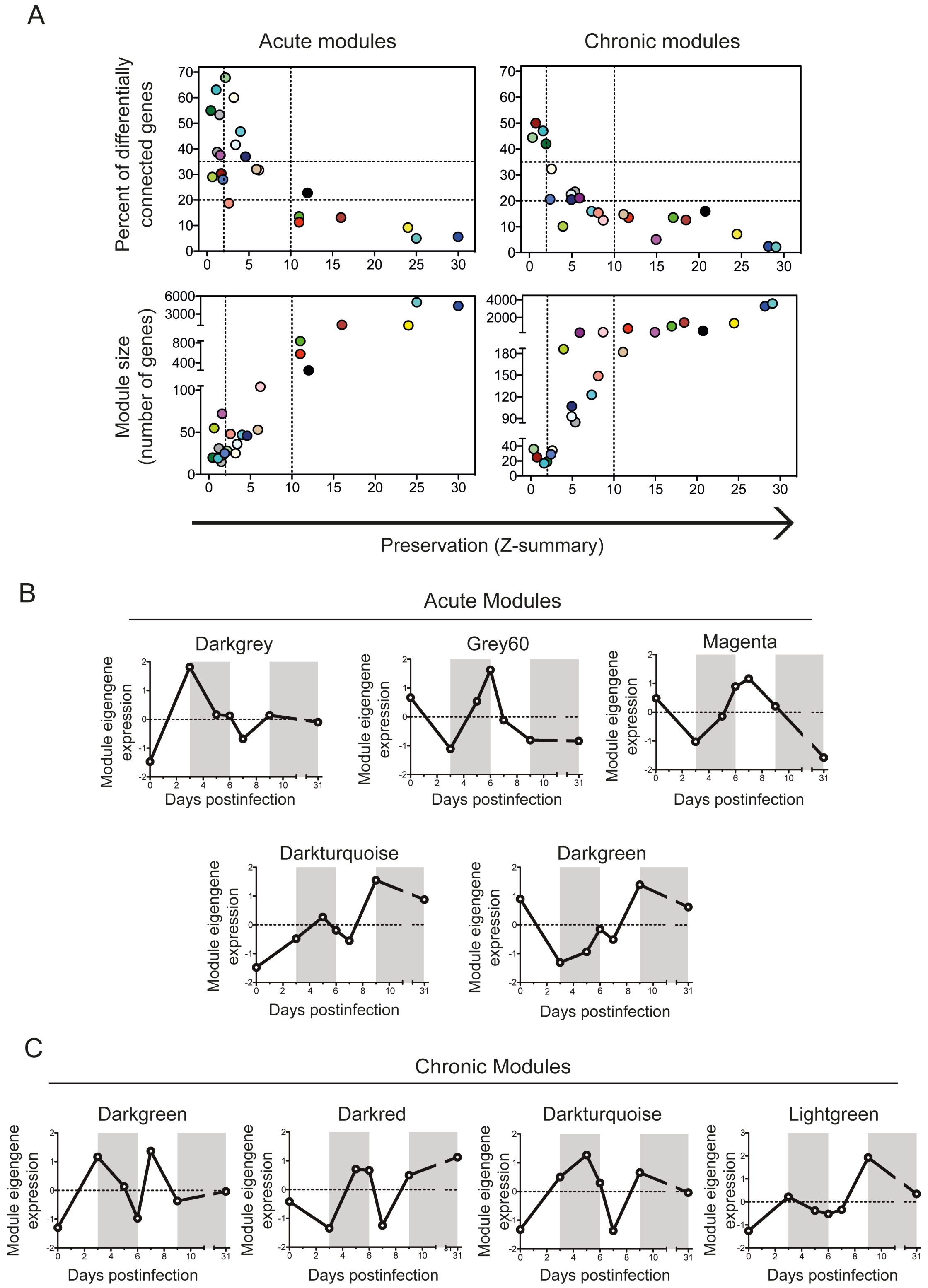


**S5 Figure. Module preservation between acute and chronic infections.** (A) Module preservation scores were compared to the percent of differentially connected genes in a module and to module size. Dashed lines indicate the thresholds (see

Materials and Methods for details). (B-C) Eigengene expression profiles of (B) acute- and (C) chronic-specific modules.

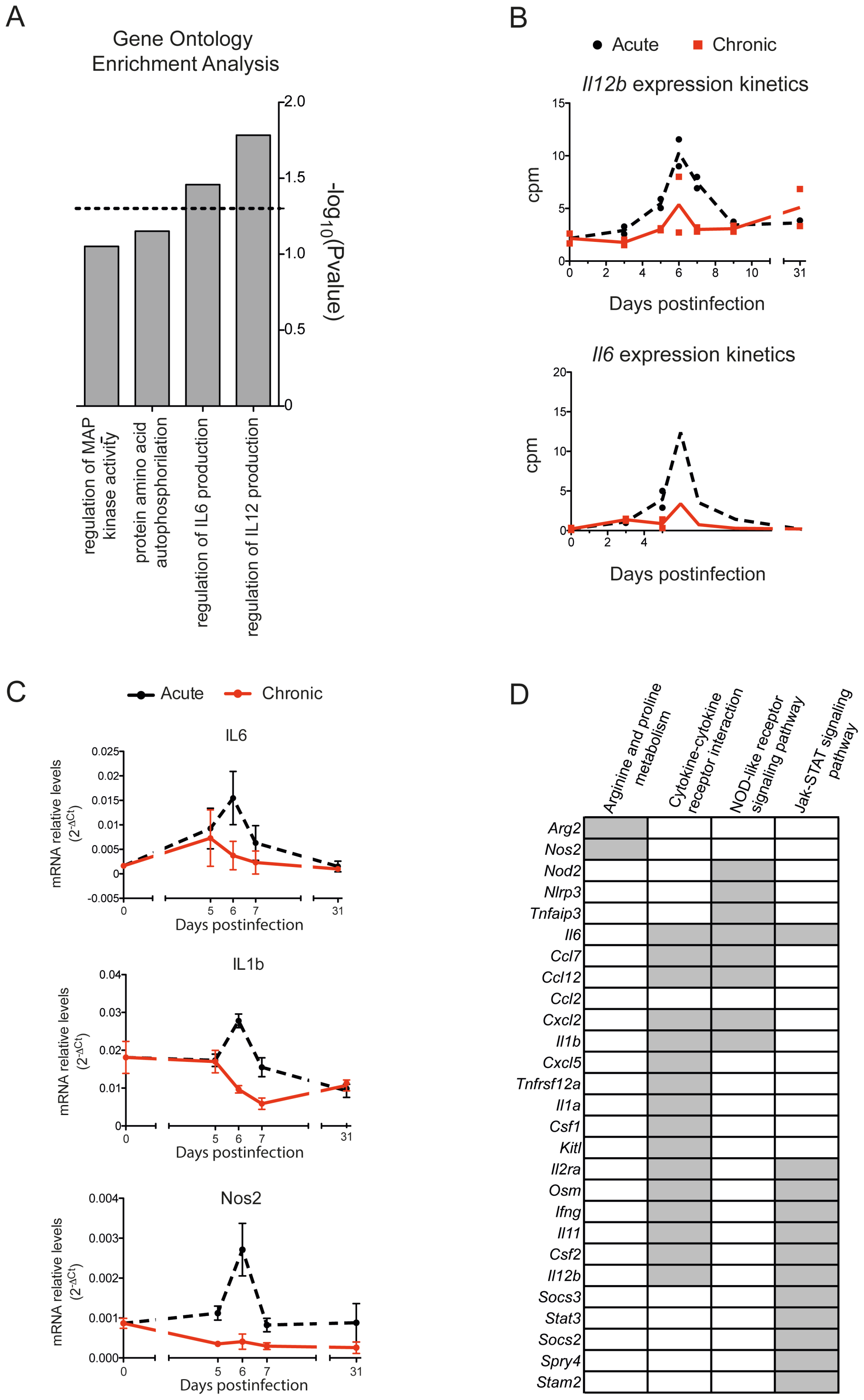


**S6 Figure. Acute-grey60-specific module unveils a differential regulation of genes related to macrophage inflammatory response in acute and chronic infections.** (A) GO terms enriched in genes from the acute-grey60 module (dashed line mark where p=0.05). (B) RNA-seq-derived *Il12b* and *Il6* expression kinetics in acute and chronic infections. (C) Quantitative PCR of *Il6*, *IL1b* and *Nos2* was performed at the indicated days postinfection from spleens of acute and chronic infected mice. Relative gene expression level was normalized by GAPDH. Data are shown as mean ± SEM from 5 to 6 mice. Significant differences were determined by an unpaired two-tailed t test. ns, non significant; * $p \leq 0.05$; ** $p \leq 0.01$. (D) List of 27 representative genes with a correlation>0.8 with *Il6* expression kinetics, and log2FC>1 of gene expression between acute and chronic infection at day 6 p.i.. Grey boxes indicate the enriched KEGG pathways that each gene belongs to.

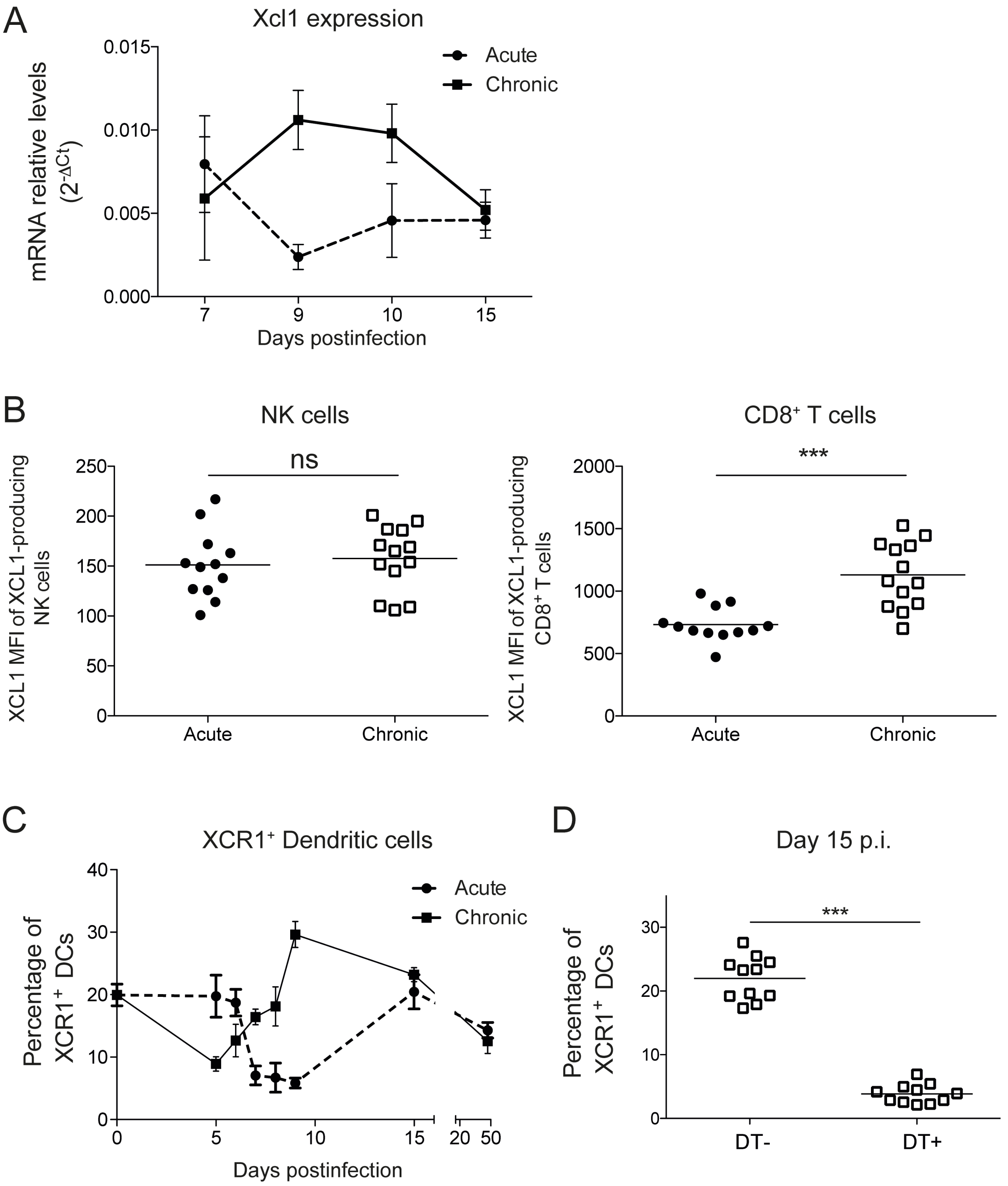


**S7 Figure. Analysis of Xcl1 expression and quantification of XCR1$^{+}$ DCs.** (A) Quantitative PCR of Xcl1 was performed at the indicated days postinfection from spleens of acute and chronic infected mice. Relative gene expression level was normalized by GAPDH. For each group and time point, the mean ± SEM is shown. (B) MFI of XLC1 in NK and CD8+ T cells. (C) Percentage of XCR1$^{+}$ dendritic cells (from CD11c$^{+}$ B220$^{-}$ cells) in spleen over the course of the infection in acute or chronic infected mice determined by flow cytometry analysis. (D) Spleen lymphocytes were harvested at day 15 postinfection from chronic infected mice non treated (DT-) or treated (DT+) with Diphtheria Toxin and percentages of $XCR1^+$ dendritic cells were assessed. Data are representative of three independent experiments. For each group the mean ± SEM of n= 11 is shown. *** $p \leq 0.001$ (Unpaired two-tailed t test).
